# Transposon mutagenesis screen in *Klebsiella pneumoniae* identifies genetic determinants required for growth in human urine and serum

**DOI:** 10.1101/2023.05.31.543172

**Authors:** Jessica Gray, Von Vergel L Torres, Emily CA Goodall, Samantha A McKeand, Danielle Scales, Christy Collins, Laura Wetherall, Zheng Jie Lian, Jack A Bryant, Matthew T Milner, Karl A Dunne, Chris Icke, Jessica L Rooke, Thamarai Schneiders, Peter A Lund, Adam F Cunningham, Jeffrey A Cole, Ian R Henderson

## Abstract

*Klebsiella pneumoniae* is a global public health concern due to the rising myriad of hypervirulent and multi-drug resistant clones both alarmingly associated with high mortality. The molecular microbial genetics underpinning these recalcitrant *K. pneumoniae* infections is unclear, coupled with the emergence of lineages resistant to nearly all present day clinically important antimicrobials. In this study, we performed a genome-wide screen in *K. pneumoniae* ECL8, a member of the endemic K2-ST375 pathotype most often reported in Asia, to define genes essential for growth in a nutrient-rich laboratory medium (Luria-Bertani medium), human urine and serum. Through transposon directed insertion-site sequencing (TraDIS), a total of 427 genes were identified as essential for growth on LB agar, whereas transposon insertions in 11 and 144 genes decreased fitness for growth in either urine or serum, respectively. These studies provide further knowledge on the genetics of this pathogen but also provide a strong impetus for discovering new antimicrobial targets to improve current therapeutic options for *K. pneumoniae* infections.

## Introduction

*Klebsiella pneumoniae* are Gram-negative bacteria found ubiquitously in natural and man-made environments (1)1} (2, 3). In immunocompromised patients, and those with co-morbidities such as diabetes or alcoholism, *K. pneumoniae* can colonize multiple body sites causing a diverse array of infections ranging from life-threatening pneumonia, urinary tract infections (UTIs), and bacteremia (4–8). *K. pneumoniae* is a frequently reported multidrug-resistant (MDR) pathogen isolated from nosocomial settings. Such infections have generally higher mortality rates and require longer treatment regimens and hospital stays, placing a significant burden on healthcare providers (9, 10). Of most concern is resistance to β-lactam antibiotics such as penicillin, aztreonam, and extended-spectrum cephalosporins, by horizontally acquired plasmid-mediated β-lactamases (11). While combined antibiotic therapies have been deployed in the clinic, the emergence of pandrug-resistant strains has created an urgency to develop new and efficient approaches to treat *K. pneumoniae* infections (12–14). The identification of genetic determinants essential for growth and those required for survival of *K. pneumoniae in vivo* may help elucidate novel therapeutic targets (15, 16).

Recent large scale phylogenetic investigations have suggested that distinct lineages of *K. pneumoniae* exist in clinical and environmental settings (17). Furthermore, two clinical pathotypes of *K. pneumoniae* have emerged: classical and hypervirulent. While the full repertoire of genetic factors required for *K. pneumoniae* virulence remains unclear, various studies have highlighted the importance of genes involved in adhesion, iron acquisition, capsulation and cell envelope biogenesis (18). Nevertheless, for both classical and hypervirulent *K. pneumoniae* strains, the ability to colonize the urinary tract and survive in the bloodstream are essential pathogenic traits and are dependent on a large and diverse range of virulence factors (19). Indeed, previous studies demonstrated that lipopolysaccharide (O-) (9 serotypes) and capsular polysaccharide (K-) antigens (79 serotypes) contribute to the resistance of serum-mediated killing and antiphagocytosis (20, 21). However, prior genetic screens have revealed that serum-resistance in Gram-negative bacteria is an intricate phenotype defined by a multitude of factors (the serum resistome) (22–26). A recent study of *K. pneumoniae* identified 93 different genes associated with serum resistance across four distinct strains but found that only three genes (*lpp*, *arnD*, and *rfaH*) played a role in all strains. These data hint that multiple mechanisms of serum resistance may exist in *Klebsiella* lineages. In contrast, the full repertoire of genes required for growth of *K. pneumoniae* in the urinary tract (the urinome) has not been defined.

To define the genetic basis of the *K. pneumoniae* urinome and serum resistome, we used Transposon Insertion Sequencing (TIS), also referred to as Transposon Directed Insertion-site Sequencing (TraDIS). TraDIS is a genome-wide screening technique that has been widely used to identify genes essential for bacterial growth and to discover conditionally essential genes under physiologically relevant conditions (27–37). In this study, we generated a highly saturated transposon library within *K. pneumoniae* ECL8, a member of the K2-ST375 phylogenetic lineage that includes hypervirulent clones of epidemiological significance known to cause infections in relatively healthy subjects predominantly in Asia (38). We identified the repertoire of genes essential for growth in laboratory conditions, in pooled human urine, and in human serum. Selected fitness-genes were validated for growth in pooled urine or serum by generating single-gene deletion mutants. Our study provides insight into the molecular mechanisms that enable *K. pneumoniae* to survive *in vivo* and cause disease.

## Methods

### Bacterial strains, plasmids, and culturing conditions

A complete list of bacterial strains, plasmids, and primers utilized in this study are listed in Table S1 and Table S2. To culture strains, scrapings from a frozen glycerol stock strain were plated onto solid Luria-Bertani (LB) (10 g tryptone, 5 g yeast extract, 10 g NaCl) supplemented with agar 1.5% (w/v) and were incubated overnight at 37°C. A single colony was isolated and used to inoculate 5 mL of liquid LB medium incubated overnight at 37°C with 180 RPM shaking. To prepare stocks, bacteria were grown to mid-exponential phase in LB medium and stored at -80°C with 25% (v/v) glycerol. When required, solid or liquid media were supplemented with appropriate antibiotics at the following concentrations: ampicillin (35 μg/mL); chloramphenicol (100 μg/mL); and kanamycin (100 μg/mL).

### Generation of the *K. pneumoniae* ECL8 transposon mutant library

An overnight culture of *K. pneumoniae* ECL8 was inoculated into 2× YT broth at OD_600_ 0.05 supplemented with a final concentration of 0.7 mM EDTA and incubated at 37°C with 180 RPM shaking. Cells were harvested at 0.4 OD_600_ by centrifugation (4,000 *g*) at 4°C for 20 min. Cell pellets were washed 4 times with ice-cold 10% (v/v) glycerol. *K. pneumoniae* ECL8 electrocompetent cells were transformed with 0.2 µL of EZ-Tn*5*™ transposon (Epicentre) at 1.4 kV and recovered in brain heart infusion (BHI) broth for 2 h at 37°C with 180 RPM shaking. Cells were plated onto LB agar supplemented with 50 µg/mL kanamycin and incubated overnight at 37°C. More than 1 million transformants were pooled in 15% (v/v) LB-glycerol for storage at -80°C until required.

### Transposon library screening in human urine

Human urine from 7 healthy male volunteers was sterilized using a vacuum filter (0.2 μM). Urine was pooled and stored at -80*°*C until required. Approximately 2 × 10^8^ *K. pneumoniae* ECL8 TraDIS mutants were inoculated into 50 mL of urine and grown for 12 h at 37*°*C with shaking (P1) in biological duplicates in parallel with control samples that were instead inoculated into 50 mL of LB medium. A sample of P1 was inoculated into 50 mL of urine or LB medium, respectively, at an initial OD_600_ of 0.05 and grown for a subsequent 12 h (P2). Both control and test experiments were passaged and grown for a further 12 h (P3). A sample of culture from P3 was removed and normalized to an OD_600_ of 1 for subsequent genomic DNA extraction.

### Transposon library screening in human serum

A 10 mL sample of human blood was collected from 8 healthy volunteers (male and female). Blood was pooled and centrifuged at 6,000 *g* for 20 min to separate sera from blood components. The pooled serum was sterilized, aliquoted and stored at -80*°*C until required. Approximately 2 x 10^8^ *K. pneumoniae* ECL8 mutants were inoculated into 1 mL of pre-warmed sera, heat-inactivated sera (60*°*C for 1 h) in biological duplicates. Samples were incubated at 37°C for 90 min with 180 RPM shaking. Following this, cells were harvested by centrifugation at 6,000 *g* for 10 min and washed twice with PBS. Cells were then inoculated into 50 mL of LB for outgrowth at 37*°*C with 180 RPM shaking and harvested at an OD_600_ of 1 by centrifugation at 6,000 *g* for 10 min. A 1 mL sample of each culture was used for genomic DNA extraction.

### Transposon library sequencing and data analysis

Pooled kanamycin resistant *K. pneumoniae* ECL8 cells were prepared for sequencing following an amended TraDIS protocol (27). Briefly, genomic DNA from the *K. pneumoniae* ECL8 library of pooled mutants was extracted from ∼1×10^9^ cells in biological duplicate, as per manufacturer’s instructions (Qiagen QIAamp DNA blood minikit). The DNA concentration of replicates was determined using the Qubit 2.0 fluorometer (Thermo Fisher Scientific). The DNA was sheared into ∼200 bp fragments by ultra-sonication (Bioruptor Plus Diagenode). Fragmented DNA samples were prepared for Illumina sequencing using the NEBNext Ultra I DNA Library Prep Kit for Illumina according to manufacturer’s instructions with the following modifications: ends of fragmented DNA were repaired using the End Prep Enzyme Mix (NEB) and an NEBNext adaptor for Illumina sequencing was ligated to the newly repaired ends. During protocol optimization, samples were analyzed using a Tapestation 2200 (High Sensitivity D5000) to determine DNA fragment sizes following adapter ligation. The uracil within the adaptor hairpin loop was enzymically excised using the USER enzyme (NEB). AMPure XP SPRI beads (Beckman Coulter) or custom-made SPRI beads were used to select for DNA fragments ∼250 bp in size.

To enrich for fragments containing the transposon a custom PCR step was introduced using a forward primer annealing to the 3’ end of the antibiotic marker of the transposon and a reverse primer that annealed to the ligated adaptor. The PCR product was purified using SPRI beads at a ratio of 0.9:1 (beads:sample). A further custom PCR step introduced sequencing flow-cell adaptors (Illumina barcodes) and inline barcodes to allow for sample multiplexing (Table S3). Samples were then stored at -20°C until sequencing. After qRT-PCR (KAPA Library Quantification Kit Illumina® Platforms) to determine TraDIS library concentration, a 1.5 μL sample of each library sample diluted to 8 nM was combined and mixed by pipetting to generate a pooled amplified library (PAL) of samples for multiplexed sequencing. The PAL was prepared for Illumina MiSeq sequencing according to manufacturer’s instructions.

Bioinformatic analysis of sequencing reads was completed on the Cloud Infrastructure for Microbial Bioinformatics (CLIMB) (39). Raw fastQ files were first demultiplexed according to Illumina barcodes automatically using the Illumina MiSeq software. Experimental replicates were separated by custom inline barcodes using the FastX barcode splitter and the barcodes were subsequently removed using the FastX trimmer (v0.0.13). Each read was checked for the presence of the transposon sequence in two steps; first, reads that contained the first 25 bp of the transposon sequence (AGCTTCAGGGTTGAGATGTGTA), introduced by the TKK_F primer, allowing for 3 mismatches were filtered and parsed. Parsed reads were subsequently checked for the presence of the final 10 bp of the transposon sequence (TAAGAGACAG), allowing for 1 mismatch. The transposon sequence and reads <20 bp were trimmed using Trimmomatic (v0.39). Resulting reads were mapped to the *K. pneumoniae* ECL8 [HF536482.1 and HF536483] genome and plasmid using the Burrows-Wheeler Alignment tool (BWA-MEM) using default parameters (i.e. -k=20bp exact match). Mapped reads were indexed using samtools (v1.8) and converted into bed format using the Bam2bed tool (Bedtools suite v2.27.1). The resulting bed file was intersected against the annotated CDS in the *K. pneumoniae* ECL8 gff file, which was generated using PROKKA (v1.14.0). Transposon insertion sites and their corresponding location were quantified using custom python scripts. Mapped transposon insertion sites were visualized using the Artemis genome browser (40).

### Statistical analysis of transposon insertion density

A modified geometric model was applied to identify the probability of finding k-1 insertion-free bases followed by a transposon insertion as reported previously (27). In a string of 10,000 independent trials the probability of an insertion ‘p’ was calculated using the following equation: P(k) = p(1-p)(k).

### Identification of putative essential genes

In-house scripts kindly provided by Langridge and colleagues (41) were amended and used for the prediction of essential genes. Briefly, the number of unique transposon insertion sites for each gene was normalized for CDS length (number of UIPs/CDS length in bp), which was denoted the insertion index score. The Freedman-Diaconis method was used to generate a histogram of the insertion index scores for each gene of the *K. pneumoniae* ECL8 TraDIS library. Distributions were fitted to the histogram using the R MASS library (v4.0.0).

An exponential distribution was applied to the ‘essential’ mode, situated on the left of the histogram. A gamma distribution was applied to the ‘non-essential’ mode, situated on the right of the histogram. The probability of a gene belonging to each mode was calculated and the ratio of these values was denoted the log-likelihood score. Genes were classified as ‘essential’ if they were 12 times more likely to be situated in the left mode than the right mode. Genes with log-likelihood scores between the upper and lower log_2_(12) threshold values of 3.6 and -3.6 respectively were classified as ‘unclear’. Genes with log-likelihood scores below the 12-fold threshold were classified as ‘non-essential’.

### Genetic context and comparison of essential gene lists between ECL8, KPNIH1, RH201207 and ATC43816

Annotated genomes of *K. pneumoniae* KPNIH1 (Accession: CP009273.1), RH201207 (Accession: FR997879.1), and ATCC 43816 (Accession: CP009208.1) were downloaded from NCBI. Nucleotide sequences of putative essential genes were extracted using the SeqKit toolbox (42). *K. pneumoniae* ECL8 essential gene homologs were identified in KPNIH1, RH201207 and ATC43816 using Galaxy AU NCBI BLAST+ wrapper blastn using the following criteria (e-value > 1e-10, percent identity >90, percent length >30) and utilizing the top hit based on calculated e value and bitscore (43).

The phylogenetic and average nucleotide identity (ANI) analysis of ECL8 and other *K. pneumoniae* isolates (Dataset S3) was performed using the Integrated Prokaryotes Genome and Pan-genome Analysis (IPGA v1.09) webserver (44). Using the default settings of the genome analysis module, ANI values between each submitted genome pairs were calculated. Then, IPGA performed genome annotation based on entries in the gcType microbial genome database for the given target genome list for phylogenetic analysis and tree construction (45).

### BioTraDIS analysis for the identification of conditionally advantageous genes

To identify conditionally essential genes, we used the Bio-Tradis analysis pipeline (16), that measures the read count log_2_ fold changes between each CDS. To ensure robust analysis, CDS with <50 sequence reads in either the test or control condition were filtered and binned. CDS with a log_2_ fold-change >2 (test/control) and a Q-value of >0.05 were classified as conferring a fitness advantage under the conditions tested. To comparatively analyze the impact of transposon insertion-events in genomic regions in the serum-resistant dataset (180 min) to the input, we used the AlbaTraDIS package (v1.0.1) with the following parameters: -a -c 10.0 -f 2 -m 0 (46).

### Generation of knock-out strains

Mutants derived from *K. pneumoniae* ECL8 were constructed using the λ Red recombinase system (47). Briefly, the *K. pneumoniae* ECL8 strain was transformed with pACBSCSE, which contains genes coding for the arabinose-induced λ Red recombinase system that permits homologous recombination between dsDNA PCR products and target loci in the bacterial genome. The recombination is based on short stretches of flanking homology arms (∼65bp) with the site of recombination. Using pKD4 as a template for the selectable kanamycin cassette, the PCR products used for replacing the target genes were amplified, gel extracted, and electroporated into electro-competent strain *K. pneumoniae* ECL8 harboring pACBSCSE prepared in the presence of 0.5% (w/v) arabinose. Candidate mutants were screened using antibiotics, verified by PCR using primers outside the region of recombination then followed by Sanger sequencing.

### Bacterial growth assay

A single colony of each bacterial strain was inoculated into 5 mL of LB medium and grown overnight as previously described. Strains were normalized to an OD_600_ of 1.00 (approximately 7 × 10^8^ cells) and washed twice in PBS by centrifugation at 6,000 *g* for 10 min. Cell pellets were resuspended in 1 mL of the growth medium required for the bacterial growth assay. Greiner Bio-One 96-well U-bottom microtiter plates were inoculated with bacterial strains at an initial OD_600_ of 0.02 in a final well medium volume of 150 μL and sealed with a Breathe-Easy® sealing membrane (Sigma Aldrich). Inoculated plates were incubated at 37°C with 300 RPM shaking, OD_600_ measurements were taken at 15 min over 24 h using a CLARIOstar® plate reader (BMG LABTECH).

### Urine co-culture competition studies between K. pneumoniae ECL8 and isogenic mutants

Overnight cultures of *K. pneumoniae* ECL8 and an isogenic mutant were normalized to an OD_600_ of 1.00 and washed twice with PBS. Cell pellets were resuspended in 1 mL of urine. A 500 μL of either the WT or mutant culture were inoculated into 25 mL of urine and incubated at 37°C with 180 RPM shaking for 12 h. The sample was then transferred to 25 mL of fresh urine at an initial OD_600_ of 0.05 and grown for a further 12 h. The culture was passaged for a further 12 h of growth. These cultures were serially diluted in PBS at 0, 6, 12, 24 and 36 h and plated onto LB agar. Following overnight growth, agar plates were replica plated onto either LB agar or LB agar supplemented with 100 μg/mL kanamycin grown overnight at 37°C then enumerated. The number of WT CFUs was calcaulated by subtracting the number of CFUs in media with antibiotics from the total number of CFUs in media without antibiotics. Relative fitness was calculated as a competitive index (CI), defined as ratio of mutant:WT viable cells divided by the corresponding inoculum.

### Lipopolysaccharide extraction

A single colony of ECL8 was inoculated into 5 mL of LB medium and incubated overnight at 37°C with 180 RPM shaking. Overnight cultures were normalized to an OD_600_ of 1.00 and centrifuged at 14,000 *g* for 10 min. Cells were resuspended in 100 μL of cracking buffer (0.125 M Tris-HCI, 4% (w/v) SDS, 20% (v/v) Glycerol, 10% (v/v) 2-mercaptoethanol in dH2O). The cell suspension was incubated at 100°C for 5 min and transferred to -80°C for 5 min. The cell suspension was incubated at 100°C for a further 5 min and centrifuged at 14,000 *g* for 10 min. An 80 μL sample of the supernatant was transferred to a new tube and incubated with 5 μL of 5 mg/mL Proteinase K (QIAGEN) for 1 h at 60°C. Samples were diluted ×2 with Laemmli sample buffer (Sigma-Aldrich) and incubated at 95°C for 5 min. Samples were loaded onto a 4-12% Bis-tris precast NuPAGE gel (Invitrogen) and run at 150 V for 1.5 h. LPS was visualized by silver staining using the SilverQuest kit (Invitrogen) following manufacturer’s instructions.

### Serum bactericidal assay (SBA)

Pooled serum was thawed on ice and pre-warmed to 37°C. Overnight cultures of bacterial strains were normalized to an OD_600_ of 1.00 in LB medium. Bacterial cells were washed twice with PBS and resuspended in a final volume of 1 mL PBS. A 50 μL sample of the cell suspension was inoculated into 50 μL of sera. Sera and cell samples were incubated at 37°C with 180 RPM shaking. Viable bacterial cells were enumerated by plating 10 μL of cell and sera suspension onto LB agar over a time-course: 0 min, 30 min, 60 min, 90 min, 180 min. Serum bactericidal activity was plotted as the log_10_ change in colony forming units (CFU) relative to the initial inoculum.

## Results and discussion

### Generation and sequencing of a *Klebsiella pneumoniae* strain ECL8 transposon input library

A *K. pneumoniae* ECL8 mini-Tn5 mutant library composed of >1 million mutants was constructed and subsequently subjected to TraDIS. The resulting reads were mapped to the reference genome that consists of a ∼5.3 Mb chromosome and a single 205-kb plasmid (EMBL Accessions: HF536482.1 and HF536483) (Figure 1A-B). The gene insertion index scores of the two technical replicates of the *K. pneumoniae* ECL8 TraDIS library (KTL1 and KTL2) were highly correlated with each other (R^2^=0.955) demonstrating a high level of reproducibility (Figure S1A). Therefore, sequence reads for both replicates were combined for downstream analysis and essential gene prediction. Through iterative rounds of sequencing, we determined that the library was sequenced to near saturation as the discovery of unique reads plateaued at ∼2 million reads (Figure S1B). This library contained 554,834 unique genome-wide transposon insertion sites and 499,919 of these insertions map within annotated coding sequence (CDS) boundaries. This high level of transposon coverage equates to an average transposon insertion every 9.93 bp throughout the genome, equivalent to an insertion approximately every four codons. A de Brujin graph visualized in the genome assembly package Bandage revealed that the sequencing coverage of the plasmid was only 1.33 fold higher than that of the chromosome indicating that the plasmid is likely present as a single copy (Figure S2) (48). Table 1 summarizes the data relating to the ECL8 libraries.

**Figure 1.**
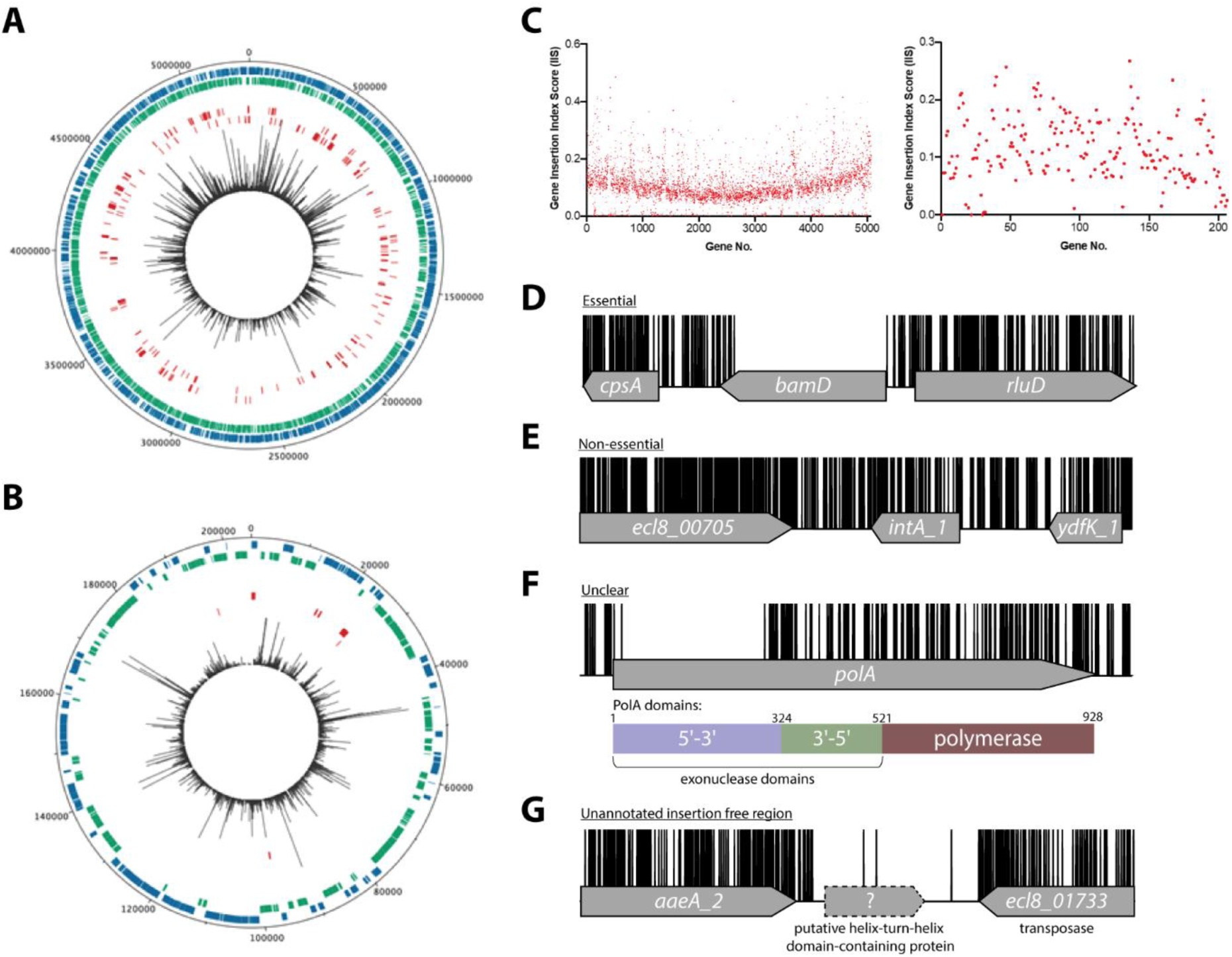
Overview of ECL8 TraDIS data: mapped insertions, insertion index profile and example gene insertion plots. Insertions are illustrated on the (A) Chromosome and (B) Plasmid, respectively. The outermost track displays the length of the ECL8 genome in base pairs. The subsequent two inner tracks correspond to coding sequences (CDS) on the sense (blue) and antisense (green) DNA strands, respectively. Putative essential CDSs are highlighted in red. The inner-most track (black) corresponds to the location and read frequency of transposon sequences mapped successfully to the *K. pneumoniae* ECL8 genome. Plot generated using DNAPlotter. (C) Gene insertion index scores (IIS) of the *K. pneumoniae* ECL8 TraDIS library mapped in order of genomic annotation of the *K. pneumoniae* ECL8 (left) chromosome and (right) plasmid. Example transposon insertion profiles categorised into essential, non-essential and unclear: (D) an essential gene – *bamD*, an essential outer membrane factor for β-barrel protein assembly; (E) a non-essential gene – *int_*A1, a redundant (several copies) integrase required for bacteriophage integration into the host genome; (F) an “unclear” gene – *polA* an essential gene in prokaryotes required for DNA replication but showed requirement for the N-terminal 5’-3’ exonuclese domain; and (G) an insertion free region suggestive of an unannotated ORF. Transposon insertion sites are illustrated in black and capped at a maximum read depth of 1.

**Table 1:** Summary of transposon-containing sequence reads and unique insertion points (UIPs) mapped to the *K. pneumoniae* ECL8.

| Sample | Mapped reads after QC | Genome-wide UIPs | CDS UIPs | Pearson Correlation of Replicates |
| --- | --- | --- | --- | --- |
| <u>ECL8 Input libraries (LB)</u> |  |  |  |  |
| KTL1 | 5,573,710 | 409,069 | 367,230 | Figure S1A |
| KTL2 | 2,854,389 | 400,915 | 361,272 |  |
| KTL (Combined) | 8,429,782 | 554,834 | 499,919 |  |
| <u>Urine output libraries</u> |  |  |  |  |
| LB | 4,324,125 | 4,324,125 | 403,381 | Figure S7 |
| Urine | 4,839,277 | 4,839,277 | 389,251 |  |
| <u>Serum output libraries</u> |  |  |  |  |
| Serum | 4,729,032 | 115,328 | 101,348 | Figure S9 |
| Heat-inactivated serum | 3,565,433 | 284,761 | 254,735 |  |
| 180 min serum exposed | 1,481,732 | 133,790 | 121,316 | Figure S10 |

### Essential chromosomal genes in *K. pneumoniae* ECL8

To classify a gene as essential or non-essential, the number of transposon insertions within each gene on the chromosome and plasmid was normalized for gene length and this value was denoted the gene insertion-index score (Figure 1C). Insertion index scores (IIS) for the *K. pneumoniae* genome followed a bimodal distribution, as has been previously described (Figure S3) (27, 41). The probability of a gene being essential was calculated by determining the likelihood of each given insertion-index score as belonging to the essential or non-essential mode. The ratio of the IIS values was denoted the log-likelihood ratio (Log_2_-LR), a metric previously used to categorize genes into essential or non-essential groupings (27, 41). To limit the number of false-positive hits identified, genes were classified as essential only if they were 12-times more likely to be within the essential, rather than the non-essential mode. Based on the above criteria, 373 chromosomal genes were assigned a log2-LR of less than -3.6 and were therefore classified as essential for growth on solid LB medium supplemented with kanamycin (Figure 1D). It should be noted that non-coding sequences such as tRNA and rRNA were excluded from this analysis. Most genes (4551) were classified as non-essential (Figure 1E). A subset of 241 genes were located between these two modes with their essentiality mode was deemed ‘unclear’ (Figure 1F). A complete list for all *K. pneumoniae* ECL8 bimodal essentiality categorization and their associated statistical significance metrics are listed in Dataset S1.

To understand the essential gene functions, we evaluated their COG (cluster of orthologous) categories; a metric that predicts functional classification against a database of known proteins. We applied a log_2_ COG enrichment index to identify COG categories that were enriched among the essential genes in contrast to the wider genome eggNOG (v5.0) orthology data (Figure S4) (49). We identified no essential genes within the COG categories for chromatin structure and dynamics (B), cell motility (N) and secondary structure (Q) suggesting genes within these categories are not required or redundant for *K. pneumoniae* growth in the utilized nutrient-rich liquid medium. However, genes involved in signal transduction (T) and inorganic ion transport/metabolism (P) were depleted, while genes involved in translation (J), cell cycle control (D) and co-enzyme metabolism (H) were the most highly enriched. This correlates with enriched COG categories shared among essential genes in other members of the Proteobacteria such as *Salmonella enterica* Typhimurium, *S. enterica* Typhi and *Caulobacter crescentus* and serves as an internal control for the validity of the library (50–52).

### “Essential” plasmid genes

The identification of eleven genes on the *K. pneumoniae* ECL8 plasmid that met the criteria to be defined as essential was an unexpected observation (Table 2). As previously noted, the bacterial copy number of the plasmid was approximated to be one. The average IIS of a gene located on the plasmid and the chromosome was comparable, 0.16 and 0.11, respectively. Due to the presence of a single plasmid per bacterial cell, and an observed even distribution of IISs for genes located on the plasmid, the lack of insertions in the 11 genes is unlikely to be due to gene dosage effects arising from the presence of multiple copies of the plasmid. Notably, the 11 plasmid-borne “essential” genes included *repB,* which is required for plasmid replication. Therefore, these genes are likely to be required for plasmid replication and stability and are unlikely to be essential for growth. This hypothesis is supported by previous observations that *K. pneumoniae* remains viable when large indigenous virulence plasmids are cured from the bacterium (53–55).

**Table 2:**
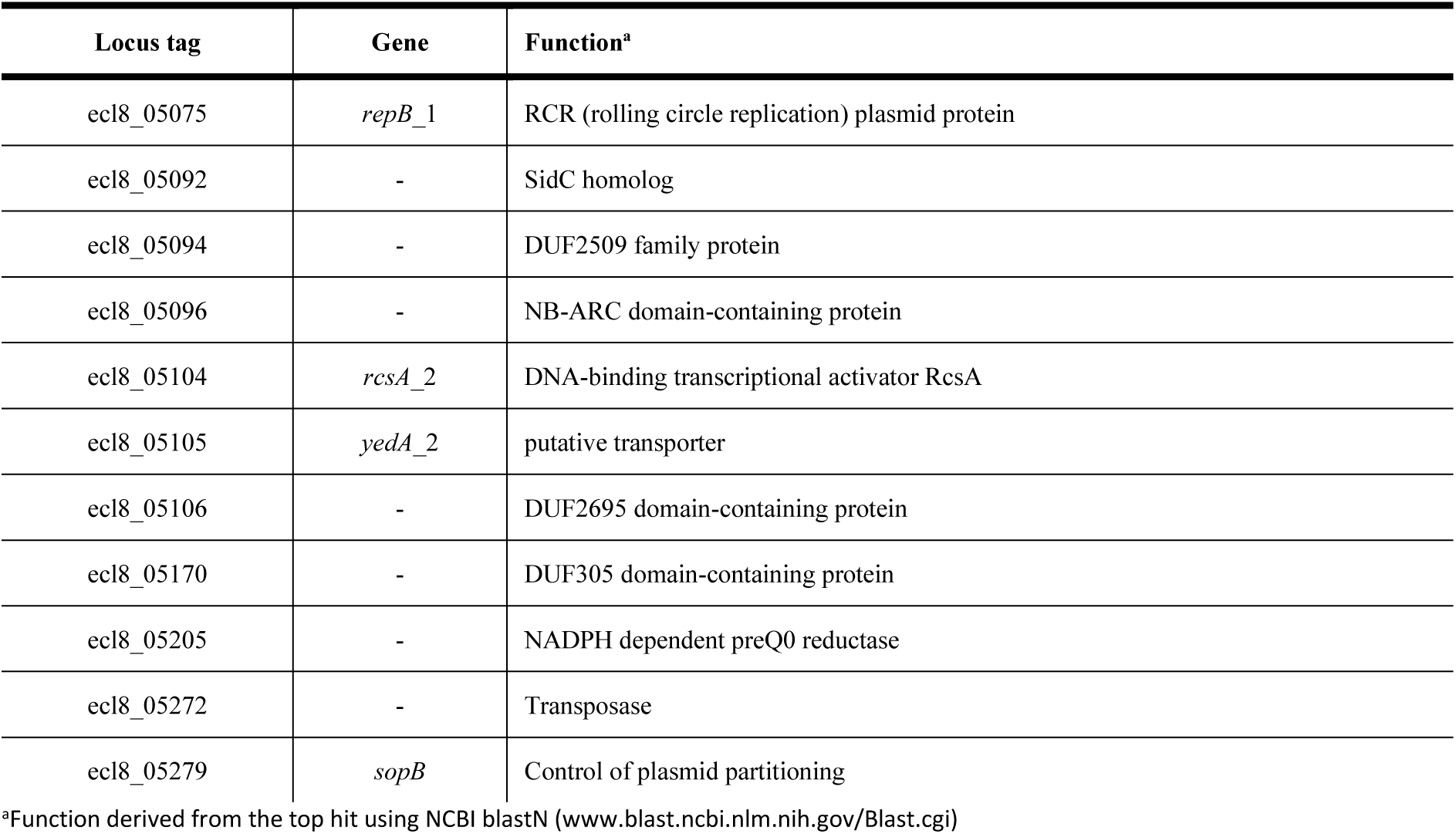
*K. pneumoniae* ECL8 plasmid-borne genes computationally deemed essential.

### Insertion free regions

Using a highly saturated transposon library it is possible to not only define essential genes, but also define chromosomal regions lacking transposons that are large enough not to have occurred by chance. Previously, we described a geometric model to predict statistically significant regions of the genome that were free from transposon insertions (27). This model indicated that with the insertion frequency calculated for this library, for a genome of 5.3 Mb, an insertion-free region (IFR) of 145 bp or greater was statistically significant (Figure S5). The size of insertion-free regions that are statistically significant for two previously published TraDIS libraries are plotted for comparison (27, 41). A total of 667 genomic regions with insertion-free regions ≥145 bp were identified, many which correspond to the 380 essential genes identified through the bimodal analysis described earlier.

The remaining insertion free regions can be explained in several ways. First, as noted previously (27), genes that encode proteins with essential domains are often excluded from the bimodal analysis described above as much of the gene contains transposon insertions. For example, the *polA* gene was classified as “unclear” but contained, barring a single transposon insertion event, an 826-bp insertion-free region. This IFR corresponds to 275 amino acids that are predicted to encode the 5’-3’ exonuclease function of the protein (Figure 1F). This region has also been shown to be required for the growth of *Escherichia coli* in laboratory conditions (56, 57). In contrast, but as noted for *E. coli polA*, multiple transposon insertions are present in the remaining portion of the gene, which confers the 3’-5’ proofreading and polymerase activity. Aside from those genes already identified by the bimodal analyses described above, our analyses revealed 54 additional genes with statistically significant IFRs, which indicates that they are likely essential for growth under the conditions tested here. Thus, our data suggest *K. pneumoniae* contains at least 427 essential genes, which we define as the ECL8 curated essential gene list.

An explanation for the remaining IFRs can be derived from traditional bioinformatic approaches to genome annotation. Originally, genome annotation protocols overlooked small ORFs because (i) they only consider genes that code for proteins >100 amino acids, and (ii) define genes based on sequence homology to those that have been previously annotated (58–60). Interestingly, the methodology used here revealed 59 regions ≥145 bp that lacked transposon insertion sites and did not map to within annotated CDS boundaries (Figure 1G). The applied approach is conservative and does not include insertion free regions that overlap with the 5’ or 3’ region of any annotated CDS. Forty-six of these regions contain potential open reading frames that could encode proteins through blastX identification. These putative unannotated ORFs were not characterized further within this study but might represent essential genes. This study emphasizes the untapped potential of TraDIS datasets to identify previously uncharacterized, essential, small coding sequences and promoter elements that are not immediately obvious by traditional and computational genomic annotation methods. A complete list of IFRs, within annotated ORFs or in intergenic regions, and an ECL8 curated essential gene list is found in Dataset S2.

### Directional insertion bias of Tn5 transposon into capsular polysaccharide (*cps*) operon

As reported previously (27, 61), due to the design of the transposon used in this study, transcription of polycistronic operons can be affected by the insertion orientation of the transposon (Figure 2A). Directional insertion bias of the transposon was noted within a large genomic region of ∼14 kb encoding the virulence-associated capsular polysaccharide (*cps)* operon (Figure 2B). The bioinformatic web-tool ‘Kaptive’ determined with high confidence the *K. pneumoniae* ECL8 *cps* operon encodes for a KL2/K2 capsule with 18/18 of the expected genes (Dataset S6) (62).

**Figure 2.**
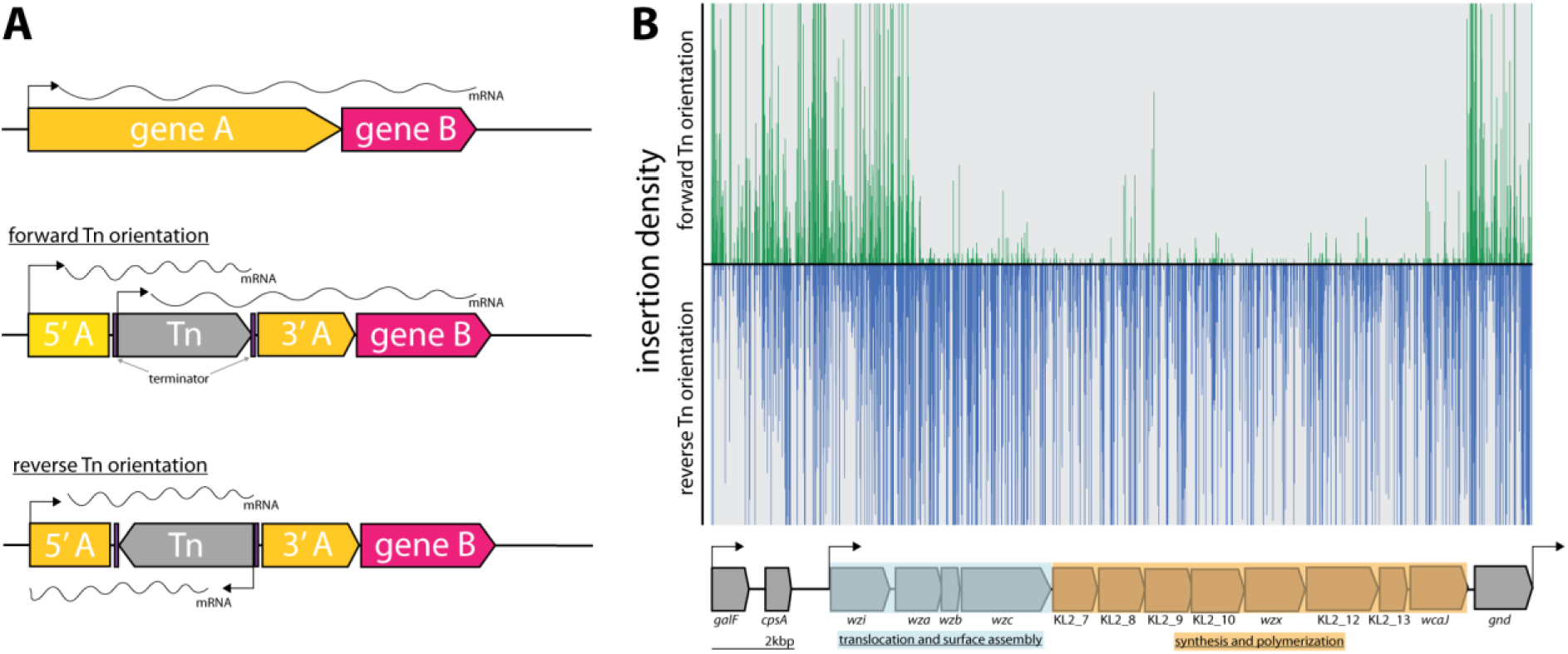
Directional insertion bias of transposon (Tn) into the *cps* operon. (A) Schematic representing the Tn orientation and effect on downstream transcription. The utilized Tn transposon is flanked by terminator sequences (purple) is shown inserted in gene A (yellow) of a hypothetical two-gene transcription unit AB in the forward or reverse orientation. In the forward orientation transcription of gene B (pink) is expected to occur from the promoters of 5’ of gene A or the internal Tn5 but polycistronic mRNA differs in length due to the attenuation by the terminators. (B) Transposon insertions mapping to the K2 capsular operon of *K. pneumoniae* ECL8. Transposon insertions are configured in the forward orientation (green), and reverse orientation (blue) and insertion densities are capped at a maximum read depth of 50. The operon structure of the K2 capsular genes consisting of three promoters driving the expression of three unidirectional polycistronic transcripts is depicted.

Notably, transposon insertions were poorly tolerated in the forward Tn orientation of the DNA strand. While bias for Tn insertions has previously been described in individual genes (e.g. ribosomal RNA genes), this study is the first to describe such a large bias across an operon (27, 50). However, the basis for this bias is not fully understood. One potential explanation for such bias might be the presence of a gene encoding a toxic small RNA (tsRNA). Typically, tsRNAs are usually present in the 5’ or 3’ untranslated regions of genes and act as regulators of gene expression at a post-transcriptional level, but due to their small size, typically <500 nucleotides, they are very difficult to predict (63, 64). However, this hypothesis does not explain why transposon insertions are uniformly absent for the opposite orientation of the transposon and not isolated to ‘hotspots’ in the 5’ and 3’ untranslated regions. An alternative hypothesis is that the reverse orientation expression of the capsular operon might result in the accumulation of intracellular capsular intermediates that may be toxic if left to accumulate in the periplasm. This finding may hint to a more cryptic regulation mechanism of the *cps* operon with implications for other similar *Klebsiella* capsular serotypes and species.

### Comparison of essential genes to other *K. pneumoniae* transposon libraries

Transposon libraries have previously been generated in other *K. pneumoniae* strains, however these studies primarily focused on identifying genes that are required for a specific phenotype i.e antibiotic stress or capsular mutants, and therefore they lack a definitive list of essential genes for *in vitro* growth (65–69). To benchmark our library, we compared our data to previously published studies that did report *K. pneumoniae* essential gene lists: KPNIH1 (ST258), RH201207 (ST258) and ATCC 43816 (ST493) (30, 70). As shown in Figure 3, the ECL8 strain is an acceptable representative for the *K. pneumoniae* KpI phylogroup as demonstrated by the phylogenetic and ANI (average nucleotide identity) analyses in the context of *K. pneumoniae* strains previously used or ‘common’ nosocomial lineages (71–74). Furthermore, our transposon mutant library is amongst the most saturated *K. pneumoniae* libraries reported to date, permitting accurate determination of gene essentiality and delineating between real and stochastic effects. The number of essential genes among the four strains ranged from 434 to 642; a complete list and comparison of our curated gene list against the reported gene lists of KPNIH1, RH201207 and ATCC 43816 is found in Dataset S3. We speculate that the higher number of essential genes predicted in the KPNIH1 gene list is due to a lower transposon density, as this is associated with an increased likelihood of false-positive results (75). This might be due in part to differences in the techniques used to construct the libraries; here we used a mini-Tn5 transposon which showed no bias in insertion whereas others have used the TnSeq method relying on the Himar I Mariner transposon, which has a requirement for a TA dinucleotide motif (76). Differences in gene essentiality might also be due to inherent genomic differences or due to differences in experimental methodology, computational approaches, or the stringency of analysis used to categorize these genes. While further validation experiments will be required to determine whether specific genes are essential, 57% of the essential genes identified in *K. pneumoniae* ECL8 were essential in the other three strains. Thus, together these data indicate that our transposon library is sufficiently dense and representative of *K. pneumoniae* (KpI phylogroup) to be used for further studies.

**Figure 3.**
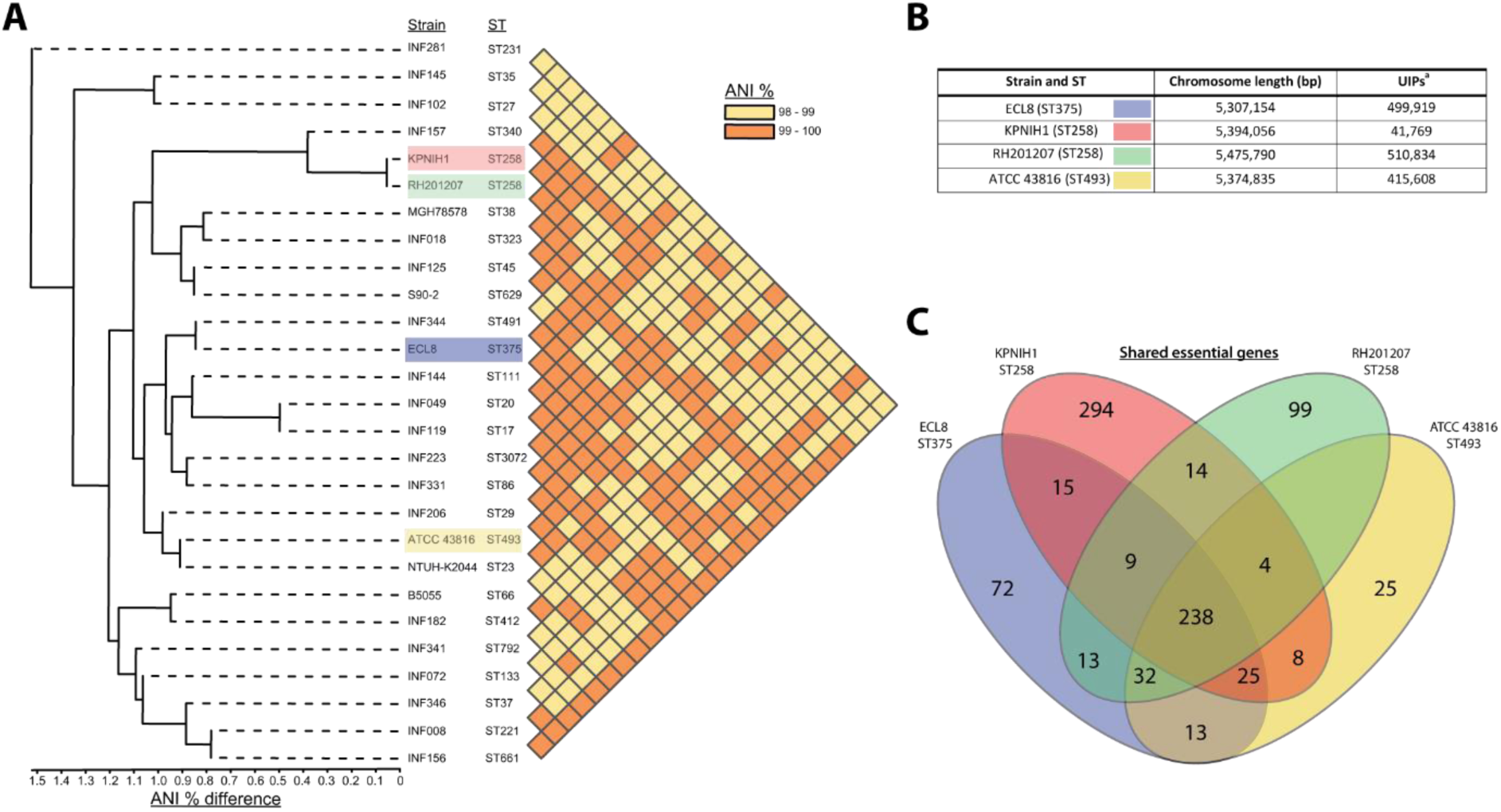
Phylogenetic context of *K. pneumoniae* ECL8 and comparison to previously reported essential gene lists. (A) Phylogenetic tree and ANI analysis generated using the IPGA webserver demonstrating the phylogenetic context of *K. pneumoniae* ECL8 compared to other previously published *K. pneumoniae* strains or isolates belonging to “21 ‘common’ lineages” of nosocomial origin by other groups. The average nucleotide identity (ANI) is a similarity index metric between a given pair of genomes applicable to prokaryotic organisms. A cutoff score of >95% indicates typically indicates they belong to the same species. (B) Tabular comparison of *K. pneumoniae* strains listing genome size and number of unique insertion points mapped (C) Venn diagram depicting the shared and unique genes required for growth in nutrient rich media (i.e. LB). Complete list of genomes, gene comparisons and exclusions lists can be found in Dataset S3.

### Identification of genes required for growth in human urine

TIS has become a formidable tool for rapid genome-scale screens to link genotype to phenotype. As the urinary tract is a major site for *K. pneumoniae* infection, we used our transposon mutant library to identify genes required for growth in urine, thus enabling us to define the urinome of *K. pneumoniae*. Cultures were grown until late stationary phase (12 h) followed by two subsequent 12 h passages before sequencing (Figure 4A). This approach eliminated mutants that were unable to grow in urine while allowing those capable of growth to flourish. DNA from test and control samples was harvested, sequenced, and analyzed as before (Figure S6). As the gene IISs for the library passaged in both LB and urine were highly correlated with respective biological replicates of 0.8483 and 0.9078 R^2^, replicates were combined for downstream analyses (Figure S7). The data for all *K. pneumoniae* genes and their associated statistical significance metrics are listed in Dataset S4.

**Figure 4.**
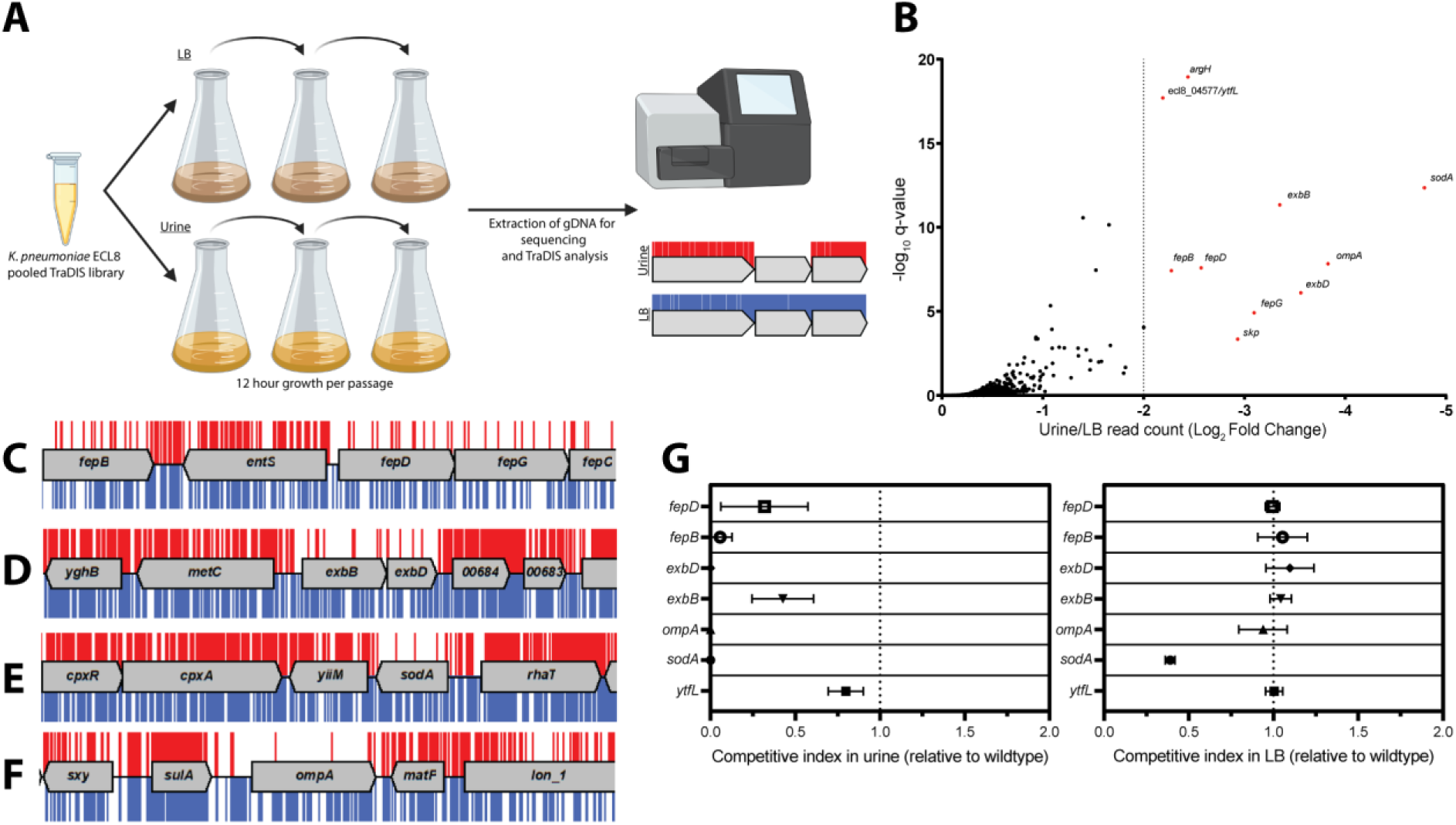
Overview and validation of ECL8 fitness-factors for growth in pooled human urine. (A) Schematic of the experimental design used to identify genes that provide a fitness advantage for *K. pneumoniae* ECL8 growth in pooled human urine. The *K. pneumoniae* ECL8 library was inoculated into either 50 mL of LB or 50 mL of urine and incubated at 37°C with 180 RPM shaking for 12 h. The library was passaged into 50 mL of fresh LB or pooled human urine at an initial OD_600_ of 0.05 two subsequent times. A 1 mL sample normalized to an OD_600_ of 1 from each culture was processed for genome extraction and multiplexed sequencing using an Illumina MiSeq. (B) Log_2_ fold change (Log_2_FC) of the read count for each *K. pneumoniae* ECL8 gene when passaged in urine relative to an LB control. Genes highlighted in red satisfy a stringent applied threshold (Log2FC >-2, Q-value ≤0.05). The Q-value is the P-value that has been adjusted for the false discovery rate for each gene. For brevity, only genes with a Log2FC ≥0 are illustrated. Selected transposon insertion profiles of genes identified as advantageous for growth in urine: (C) *fepB, fepD, fepG* (D) *exbB, exbD* (E) *sodA* and (F) *ompA*. These genes exhibited a significant loss of transposon insertions following growth in urine (red) in comparison to LB broth (blue). A 5-kb genomic region including the gene is illustrated. Reads are capped at a maximum depth of 1. (G) The fitness of gene replacement mutants relative to WT *K. pneumoniae* ECL8 in either LB medium or urine. The relative competitive index of single-gene replacement mutants after 12 h passages ×3 in either LB medium or urine. A relative fitness of one would indicate comparable fitness to WT. The mean is plotted (± 1 SD).

Ten genes that satisfied a stringent threshold of a log_2_ fold change (log_2_FC) >-2 and a Q-value ≤0.05, relative to the LB control, were considered as advantageous for *K. pneumoniae* ECL8 growth in urine (Figure 4B). The transposon insertion profiles of these genes showed a marked decrease in overall insertions following passaging in urine when compared to the LB control (Figure 4C-F), suggesting loss of these genes confers a fitness defect. These included genes encoding the superoxide dismutase SodA, the structural outer membrane protein OmpA (63), the periplasmic chaperone Skp (77), and five proteins (*fepB*, *fepD*, *fepG*, *exbB* and *exbD*) associated with iron acquisition (78). The remaining two genes (*ytfL* and *argH*) play key roles in controlling cytoplasmic cadaverine and putrescine concentrations and amino acid biosynthesis, respectively (79, 80). The proper regulation of cadaverine and putrescine maintains intracellular pH concentrations and reduce oxidative damage to proteins and DNA and may be in response to acidic conditions in urine known to be unconducive for colonization of uropathogens (81, 82).

To experimentally validate our observations, seven independent mutants were constructed by allelic replacement of the chromosomal gene with a kanamycin (*aph*) cassette. To determine whether *K. pneumoniae* strains lacking the genes identified in this screen were truly less fit in urine compared to the parent strain, the WT ECL8 and individual isogenic mutant strains were inoculated into urine or LB medium at a 1:1 ratio. These cultures were sequentially passaged, and the CFU/mL of both the WT and mutant strain were determined over a time-course of three 12 h passages (Figure 4G). When compared to the wild-type, a Δ*sodA*::*aph* mutant was less fit in both urine and LB, but the phenotype was exacerbated in urine. In contrast, the fitness of the remaining mutants was comparable to the parent strain when grown in LB medium and only less fit when grown in urine. While the relative competitive index of the Δ*ytfL*::*aph* strain was 0.8 in urine, the *ompA*, *fepB*, *fepD*, *exbB* and *exbD* mutants were dramatically less fit than the parent strain in urine (Figure 4G).

Based on these observations, we hypothesized that iron-levels were limiting growth in human urine and that the ability to acquire iron was crucial for *K. pneumoniae* to survive and grow *in vivo* (83, 84). To test this hypothesis, urine was supplemented with exogenous ferrous iron (100 μM FeSO_4_) or an iron chelator (100 μM bipyridine/2,2-dipyridyl). No variance in optical density was observed after passaging in LB medium (Figure 5A). However, the optical density of *K. pneumoniae* cultures decreased after passage in urine (Figure 5B), and the restricted growth was further exacerbated by the addition of the iron chelator 2,2-dipyridyl (Figure 5C). In contrast, supplementation of urine with exogenous iron increased growth relative to non-supplemented cultures with no difference in growth upon passage (Figure 5D). To further interrogate the role of iron limitation, the growth kinetics of the constructed mutants were grown in urine with and without exogenous ferrous iron for 16 h in a microtiter plate assay. Not unexpectedly, area under the curve (AUC) analysis revealed that mutants lacking genes involved in iron uptake (*fepB*, *fepD*, *exbB*, *exbD*) were drastically less fit than the wildtype (Figure 6E). Surprisingly, our screen showed several siderophore synthesis genes did not confer a statistically significant fitness advantage for growth in urine (i.e. *entACDF*, Figure 5F). In contrast, the highest Log_2_FC increase (∼7-fold) in read counts mapped to the <u>F</u>erric <u>U</u>ptake <u>R</u>egulator (*fur*) gene indicating its loss is beneficial for growth in urine. In Gram-negative bacteria, Fur is a key regulator for iron homeostasis functioning as a transcriptional repressor for siderophore synthesis genes in an iron concentration-dependent manner (85–87). Furthermore, the Fur regulon includes the cognate ferric enterobactin uptake (Fep) genes, for which decreased Log_2_FC read counts were also noted (Figure 5G). Altogether, this suggests that the Fur regulon plays a pivotal role for *K. pneumoniae* growth in urine by tightly regulating enterobactin synthesis and enterobactin-mediated iron uptake.

**Figure 5.**
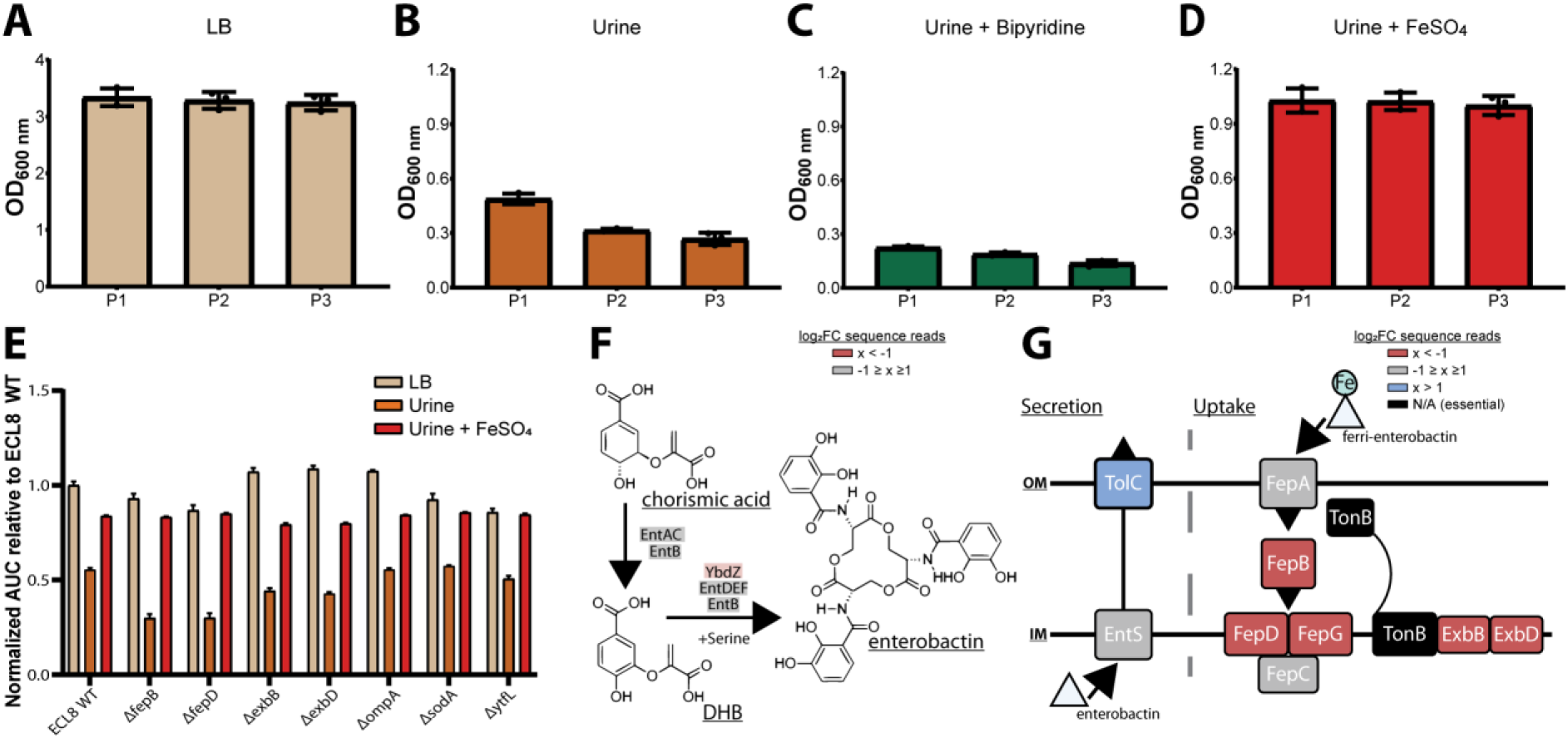
Growth of the *K. pneumoniae* TraDIS library following passaging in LB and urine and schematic diagrams of enterobactin synthesis, secretion and uptake. The OD_600_ of the *K. pneumoniae* TraDIS library following 12 h of growth (P1) and two sequential 12 h passages (P2 and P3). The library was passaged into fresh medium (A) LB or (B) urine to an initial OD_600_ of 0.05. To determine the effect of iron supplementation and depletion, urine was supplemented with exogenous iron (C) 100 μM FeSO_4_ or an iron chelator (D) 100 μM 2,2-dipyridyl. The average OD_600_ of three biological replicates for each time point is plotted (±) 1 SD. (E) Area under curve comparative analysis (OD_600_ vs. time) of *K. pneumoniae ECL8* and mutants grown in LB, urine or urine supplemented with 100 μM FeSO_4_ grown for 16 hours with 180 rpm shaking. The average of three biological replicates is plotted for each condition with error bars representing SD. (F) Simplified schematic of the enterobactin synthesis pathway. YbdZ, a co-factor of EntF for the terminal steps for enterobactin synthesis, depicted in light red had a Log_2_FC sequence read value of -1.58 suggesting this gene conferred an overall fitness advantage for growth in urine. (G) Schematic representation of enterobactin secretion and uptake. The TonB transport system is present in Gram-negative bacteria and is required to transport Fe-bound enterobactin through the outer (OM) and inner membrane (IM) to the cytosol where it can be utilized. Based on Log_2_FC sequence read value, loss of TolC (blue) was beneficial for growth, relative to an LB control. Loss of proteins, colored in red, had Log_2_FC sequence read values <-2 suggesting they confer a fitness advantage for grown in urine. Proteins depicted in grey were genes that had Log_2_FC values that ranged from -1 to 1 exposed to urine relative to an LB control. Genes depicted in black were essential and had no determinable Log_2_FC value.

**Figure 6.**
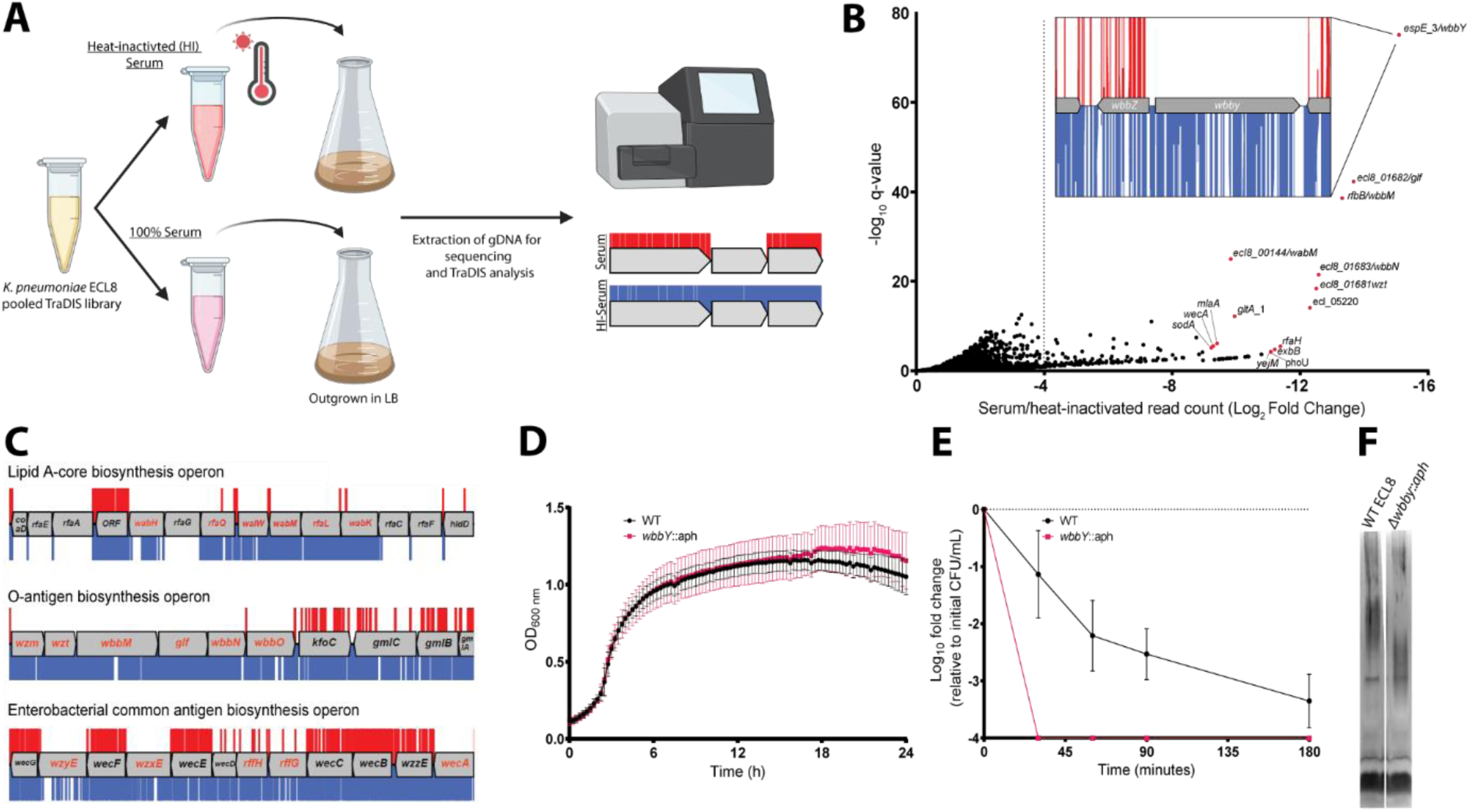
Overview and validation of ECL8 genes that increase resistance to complement-mediated killing. (A) The experimental methodology utilized for screening the TraDIS library in human serum and a heat-inactivated serum control. *K. pneumoniae* ECL8 (2x10^8^ cells) of the mutant library was inoculated into either 1 mL of human serum or 1 mL of heat-inactivated human serum and incubated for 90 min. Following exposure to serum, cells were grown to an OD_600_ of 1 in LB medium to enrich for viable mutants. A 1 mL sample normalized to an OD_600_ of 1 from each culture was processed for genome extraction and multiplexed sequencing using an Illumina MiSeq. (B) Log_2_FC for each gene of the *K. pneumoniae* ECL8 TraDIS library when incubated in pooled human serum relative to a heat-inactivated serum control. Selected genes highlighted in red are amongst the total of 144 genes that satisfy a stringent applied threshold (Log_2_FC ≥-4, Q-value ≤0.05). For brevity, only genes with a Log_2_FC ≥0 are illustrated. Inset: transposon insertion profile of *wbbY*, gene with the highest fold Log_2_FC, flanked by *wbbZ* and a transposable element at its 3’. Transposon insertions following exposure to serum and a heat-inactivated serum control are illustrated in red and blue, respectively. Transposon reads have been capped at a maximum of 10. (C) Transposon insertion profiles of genes within the: LPS, O-antigen and the ECA biosynthesis operons. Genes in red font had a significantly (Log_2_FC = ≥-4, Q-Value = ≤0.05) decreased fitness when disrupted with a transposon following exposure to serum for 90 minute (red), relative to a heat-inactivated serum control (blue). Operons are not drawn to scale and reads capped at a maximum read depth of 1. (D) Growth profile of WT *K. pneumoniae* ECL8 and Δ*wbby*::*aph* in LB broth. Mean is plotted (± 1 SD). (E) Serum killing assay of WT *K. pneumoniae* ECL8 and Δ*wbby*::*aph.* Mean is plotted (± 1 SD). (F) LPS profiles of WT *K. pneumoniae* ECL8 and Δ*wbby*::*aph*. Overnight cultures of each strain were normalised to an OD_600_ of 1. The LPS was separated on 4-12% Bis-Tris gels and was visualized by silver staining using the SilverQuest kit (Invitrogen).

### Identification of genes required for resistance to complement-mediated killing

To identify genes that protect against complement-mediated killing,previously defined as the serum resistome (22, 23), we first established the serum killing kinetics of *K. pneumoniae* ECL8. Approximately, 2 x 10^8^ cells were incubated with 100 μL of human serum for 180 min. At specified time points, aliquots were plated to determine the number of viable bacteria. As expected, no viable bacteria were recovered from the *E. coli* BW25113 control (88). In contrast, a 2 log_10_ decrease in the number of viable *K. pneumoniae* cells was observed after 90 min exposure to normal human serum, and a >4 log_10_ decrease after 180 min exposure. To determine whether the observed reduction in viable cells was due to the presence of active complement proteins, *K. pneumoniae* was incubated inheat-inactivated serum control where no reduction in viable was noted (Figure S8).

Subsequently, 2 x 10^8^ cells from the ECL8 TraDIS library were inoculated into 1 mL of human serum or 1 mL of heat-inactivated human serum and incubated for 90 min. Following exposure to serum, cells were grown in duplicate to an OD_600_ of 1 in LB medium to enrich for viable mutants. DNA was harvested from each sample, and subsequently sequenced as described previously (Figure 6A). The Pearson correlation coefficient (R^2^) of gene insertion index scores (IIS) for two sequenced biological replicates were calculated as 0.91 (normal human serum) and 0.76 (heat inactivated serum) (Figure S9). To ensure robust data analysis, thresholds were applied to the data where genes with less than 50 mapped reads in the input library were removed from downstream analysis to exclude confounding essential genes and minimize the effect of stochastic mutant loss. A total of 356 genes with a log_2_FC ≤-2, P<0.05, and Q-value of ≤0.05 between the serum and heat-inactivated control were classified as genes that are advantageous for growth under complement-mediated killing conditions (Figure 6B). The complete list of all genes analyzed for growth in human serum and their associated statistical significance metrics are listed in Dataset S5. The operons encoding lipopolysaccharide (LPS) core antigen, O-antigen, and enterobacterial common antigen (ECA) biosynthesis were noted as containing several conditionally essential genes (Figure 6C). Mutants defective in these cell envelope structures have previously been reported to play a role in membrane integrity and resistance to serum-killing for a variety of Gram-negative bacteria (89–91). Genes implicated as important for serum resistance encode proteins with an important role in membrane stability. They include GalE involved in LPS sugar sub-unit biosynthesis (92, 93), outer membrane protein MlaA involved in phospholipid retrograde transport (94), cardiolipin synthase ClsA (95), Bam complex component BamE (96), and proteins involved in LPS biosynthesis (LpxC) and its regulator (YejM) (97).

To validate our observations, we selected the gene with the greatest Log_2_FC decrease between serum and a heat-inactivated control; a putative glycosyltransferase *espE*_3/*wbbY* (inset, Figure 6B). Analyses using the ‘Kaptive’ webtool suggested that ECL8 encodes the O-antigen serotype O1v2 with the operon having 98.98% nucleotide identity to the reference genome (Dataset S6) (62). The O-antigen of the O1v2 serotype features distal repeating D-galactan I and D-galactan II sugar sub-units (98). Despite its role in O-antigen biosynthesis, *wbbY* is not located within the *rfb* LPS operon. To determine the role of *wbbY* in serum sensitivity, the gene was replaced with an *aph* cassette and the resulting mutant was compared to its parent strain using a serum bactericidal assay. A comparable growth profile was observed for *K. pneumoniae* ECL8 and the Δ*wbbY*::*aph* mutant in LB medium (Figure 6D). However, the Δ*wbbY*::*aph* mutant was more sensitive to human serum than the parent with no viable mutants surviving after 30 min (Figure 6E). To investigate whether the *wbbY* deletion affected LPS production, crude LPS extracts were visualized using silver staining (Figure 6F). The Δ*wbbY*::*aph* strain produced LPS with a lower molecular-weight compared to the parent strain, suggesting that a truncated O-antigen was expressed Alterations in the amount or length of LPS produced are known to influence susceptibility to serum mediated killing (90). Importantly, Hsieh *et al* have previously reported that deletion by *wbbY* in *K. pneumoniae* NTUH-K2044 (ST23) resulted in LPS profile defects due to abrogated D-galactan II production (98). The authors also demonstrate that gene complementation restores LPS profiles and serum-resistance, thus demonstrating that this screen was robust in identifying a key *K. pneumoniae* gene involved in complemented-mediated killing.

### Identification of genes required for serum-resistance

Following the identification of genes that when mutated confer increased sensitivity to complement-mediated killing (i.e. the serum resistome), the *K. pneumoniae* ECL8 TraDIS library was screened for mutants that confer increased resistance to serum-killing. Approximately 2 x 10^8^ TraDIS mutants were incubated in 1 mL of human serum for 180 min. The output pool was washed with PBS and plated onto LB agar supplemented with kanamycin (Figure 7A). Following overnight growth, ∼150,000 colonies were recovered and pooled for sequencing confirming good correlation of replicates (Figure S10). We then used AlbaTraDIS to identify mutations that conferred a fitness advantage or disadvantage for survival in serum. In addition to identifying genes with an increase or decrease in reads between the condition and control samples, AlbaTraDIS will also analyze changes in sequencing read depth of insertions immediately upstream and downstream of a gene, as well as within-gene transposon orientation biases. Such differences can be strong indicators of mutations that result in gene expression or regulatory effects (For a detailed explanation please see (46)). The data for insertions with a significant (>2-fold) change in read-depth following serum exposure for 180 min are listed in Dataset S7. As expected, similar to our previous experiment genes with the greatest significance in decreased insertions, suggesting decreased fitness, were O-antigen outer-core biosynthesis gene espJ_2/*wabI* and O-antigen ligase rfaL/*waaL* (Figure 7B). Genes with significantly increased insertions, suggesting increased fitness, included *hns*, *gnd* and *igaA* (Figure 7C). Closer inspection of the transposon insertion profiles further confirmed the enrichment of *hns* mutants when comparing the serum exposed test to the input control (Figure 7D). To validate our observations, a defined *hns* mutant was constructed by allelic replacement with an *aph* cassette in the same orientation as native gene transcription. A comparable growth profile in LB was observed for the *hns* mutant relative to the parent (Figure 7E). To determine whether the *hns* mutant was indeed more resistant to serum a sample of 1 x 10^8^ cells was incubated in serum for 360 min. At 30 min post-incubation the number of viable *hns*::aph cells was decreased by ∼2 log. However, no additional decrease in viable cell numbers was observed at later time points and these strains proliferated in serum (Figure 7F). Ares and colleagues demonstrated that *rcsA; galF; wzi;* and *manC* were derepressed in a Δ*hns K. pneumoniae* mutant resulting in increased capsule production (99). Similarly, up-regulation of *rcsA* was noted in an *E. coli* Δ*hns* mutant. RcsA is known to positively regulate the capsular locus (100, 101). Notably, IgaA represses the Rcs regulon, preventing activation of RcsA. Mutants in *igaA* are enriched in our experiments (Figure S11A). These data suggest that activation of the Rcs regulon confers resistance to serum killing in *K. pneumoniae*, as noted for other members of the *Enterobacteriaceae*, by increasing capsule production (102).

**Figure 7.**
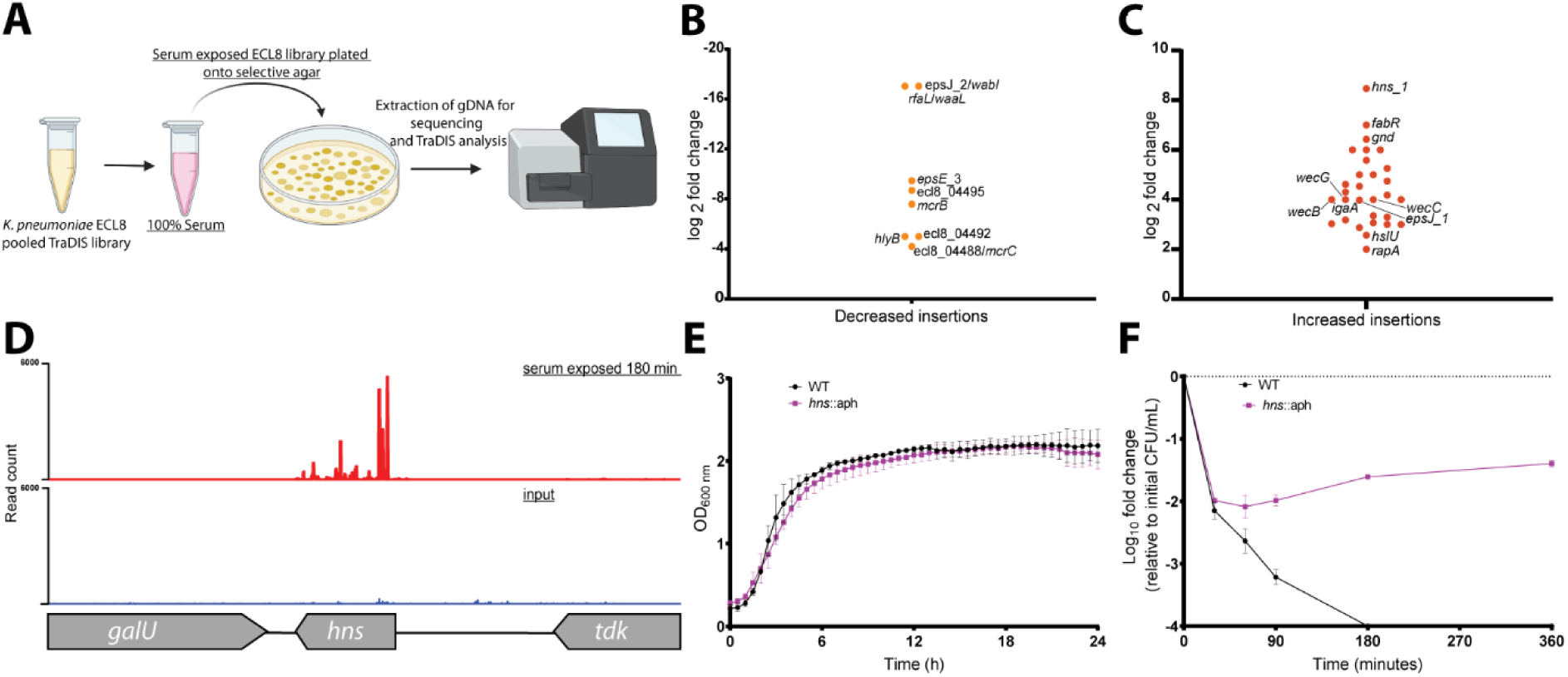
Overview and validation of ECL8 fitness-factors for survival in pooled human serum. (A) The experimental methodology utilized for screening the TraDIS library to identify genetic factors that increase resistance to human serum. *K. pneumoniae* ECL8 (2×10^8^ cells) of the mutant library was inoculated into 1 mL of human serum and incubated for 180 min and compared to before serum exposure input control. The output pool was washed with PBS and plated onto LB agar supplemented with kanamycin. Following overnight growth, ∼150,000 colonies were recovered and pooled for sequencing. Comparative analysis using AlbaTraDIS software depicting genes with (B) decreased insertions suggesting a loss of fitness or (C) increased insertions suggesting a gain of fitness to serum exposure. (D) Transposon and read count insertion profiles of *hns* locus: red illustrating pooled mutant serum exposed for 180 min and blue denoting the before serum exposure input control. (E) Growth profile of WT K. pneumoniae ECL8 and Δ*hns*::*aph* in LB broth. Mean is plotted (± 1 SD), where n=3. (F) Serum killing assay of WT *K. pneumoniae ECL8* and Δ*hns*::*aph*. Mean is plotted (± 1 SD), where n=3.

The *gnd* gene encodes a 6-phosphogluconate dehydrogenase involved in the oxidative branch of the pentose phosphate pathway. Interestingly, sequence reads mapping to *gnd* were enriched for the forward orientation of the transposon only (Figure S11B). Transposon insertions in this orientation have transcriptional read-through driving expression of the downstream neighboring gene *manC* and more distal O-antigen biosynthesis genes (i.e. *rfb* cluster) (103). The *manC* gene encodes mannose-1-phosphate guanylyltransferase that is required for the synthesis of capsule, and as noted above, upregulation of *manC* results in increased production of capsule.

## Conclusion

For a variety of reasons, the determination of gene essentiality from TraDIS data is challenging (27). Here, we used two different computational approaches to identify essential genes ensuring that our essential gene dataset includes those genes that can tolerate insertions in regions of the gene but where specific domains are essential for growth. Coupled with the presented phylogenetic and genomic analyses, the observation that 57% of the essential genes identified in this study were found to be essential in three of the four *K. pneumoniae* TraDIS libraries described to date provides reassurance that this library would provide meaningful data for subsequent studies. A minor caveat that is worthwhile mentioning is the potential for false-positive gene essentiality calls due to nucleoid-assocaited proteins (NAPs) hinders traposon insertion (104). Nevertheless, the high density of insertion sites achieved in our study ensured reliable identification of essential and conditionally-essential genes and a low probability of false positive identification, which had previously been noted for low complexity libraries (75).

The search for genetic determinants of bacterial survival and growth *in vivo* has been augmented by whole genome approaches to infection, including transposon insertion site sequencing techniques such as TraDIS and TnSeq (15). These data have potential implications for the design and development of novel drugs, vaccines and antivirulence strategies. In this study, we employed TraDIS to identify genes which, when mutated, confer decreased or increased fitness for growth in human urine and serum, two *in vivo* niches relevant to *K. pneumoniae* infection. Unexpectedly, our study revealed only 11 genes significantly required for growth in urine. These included genes encoding the outer membrane protein OmpA, its chaperone Skp, the superoxide dismutase SodA, and genes required for iron acquisition by the enterobactin uptake system. Notably, not all the genes for enterobactin synthesis, export and uptake were essential (Fig. 4). The gene encoding FepB, the periplasmic chaperone for the iron laden siderophore, was essential, as were *fepD* and *fepG*, which encode the inner membrane uptake system. The TonB complex (TonB-ExbB-ExbD), which energizes the translocation of iron laden enterobactin through the FepA outer membrane pore, was also essential for growth in urine. In contrast, the genes encoding FepA, the inner membrane ATPase FepC, and the enterobactin synthesis (Ent) and export (TolC) components were not essential. These observations can be explained in two ways. First, as *K. pneumoniae* ECL8 contains paralogous copies of *fepA* (*fepA*_1 and *fepA*_2) and *fepC* (*fepC*_1 and *fepC*_2), the non-essential nature of *fepA* and *fepC* might be the result of functional redundancy. Second, as TraDIS is in principle a large-scale competition experiment between thousands of mutants, mutants lacking enterobactin (or aerobactin) synthesis and export genes can “cheat” by acquiring the siderophore released from siderophore-producing strains within the population, a phenomenon described previously (105, 106). It should be mentioned, our study utilized filter sterilized urine which may preclude identification of genes that are required for interspecies predation, cross-protection and cross-feeding in the context of a urinary microbiome (107, 108).

In contrast to the limited number of genes in the urinome, the serum resistome consisted of more than 144 genes. This number is significantly greater than reports of a recent study that defined the serum resistome of four independent *K. pneumoniae* strains (22). While that study found 93 genes that were required for survival in one or more strains, remarkably only three genes were common to all four strains: *rfaH*, *lpp* and *arnD*. Comparison with our dataset revealed that in contrast to *rfaH, arnD* was not required for serum resistance in *K. pneumoniae* ECL8. Surprisingly, comparison of our data with the 56 genes comprising the serum resistome of the more distantly related multidrug resistant *E. coli* strain EC958 revealed 14 genes in common. The variation in the outcome of these experiments can be accounted for in different ways. First, in all cases the genes encoding O-antigen and/or capsule synthesis were required for growth in normal human serum; As the genes encoding these surface structures vary with serotype, they are not identified as common genes, though ostensibly they have the same function Second, the methodology appears at first glance to be identical in each study (growth in the presence of serum for 90 min). However, not all strains exhibit the same killing kinetics in normal human serum. Using fixed timepoints, strains with longer killing kinetics are likely to have fewer genes identified in TraDIS experiments, whereas strains with short killing windows will have more. Third, several of the genes identified in the serum resistome of other strains were excluded from our downstream analysis as there were fewer than 50 transposon insertions in the gene following exposure to heat-inactivated serum. A case in point is the *lpp* gene. This observation indicates that *lpp* is required for *K. pneumoniae* ECL8 growth in human serum but not resistance to complement dependent killing *per se*. Fourth, the density of the library used, the level of comparative annotation, and the thresholds to define significance, all have the potential to influence the final outcome of such experiments. Finally, strain-specific features such as gene duplication events, functional redundancy in biosynthetic pathways, and the variation in the genetic complement of each strain, are likely to influence the contribution each gene makes to serum resistance.

Despite the intra and interspecies variation in the serum resistome noted above, comparison of the urinome and serum resistome of *K. pneumoniae* ECL8 revealed that mutation of five genes (*ompA*, *galE*, *exbB*, *exbD* and *sodA*) conferred reduced fitness for both growth in urine and survival in serum. The role of *exbB* and *exbD* are described above but their detection in both experimental conditions reinforces the importance of iron acquisition for infection. The major outer membrane protein OmpA identified in this study encodes a functionally pleiotropic outer membrane protein (85, 86). Notably, a role for OmpA in serum resistance has previously been described for *E. coli*, however it was not identified by TraDIS experiments investigating the serum resistome (23). The molecular basis for this discrepancy remains unresolved. GalE is known to play a key role in serum resistance for *E. coli* and *Salmonella* spp.. The gene *galE* encodes a UDP-glucose 4-epimerase catalyzing the interconversion of UDP-D-galactose/UDP-D-glucose (92). It has been previously reported that GalE plays a role in D-galactan II synthesis, an O1 LPS component in *K. pneumoniae* (93, 109). These data suggest that loss of *galE* results in a modified LPS in *K. pneumoniae* resulting in increased serum susceptibility. Additionally, the manganese-dependent superoxide dismutase SodA converts superoxide to hydrogen peroxide and oxygen to prevent DNA and other cellular damage (110, 111). A previous study in *K. pneumoniae* has reported the inducible expression of *sodA* in a triumvirate system with *sodB* and *sodC*, expressed in response to anaerobic challenge and high-oxygen concentration (112). These transcriptional responses might very well be present in both the microenvironments of blood vessels and the bladder, in the form of nutrient limitation and oxygen exposure (113–115). Hence, the observed importance of *sodA* might be a consequence of elevated levels of reactive oxygen species (ROS) due to iron-acquisition and could represent a specialized metabolic adaptation for growth in these environments. Ultimately, these findings underline the interplay between bacterial iron homeostasis and oxidative stress previously reported in Gram-negative bacteria (116–120). However, these data shine a light on cross-niche (i.e. urine and serum) genes and associated pathways as suitable drug development targets to treat *Klebsiella* infections. Finally, this library was then used to identify genes conferring serum resistance. An interesting finding was the observed transposon insertion orientation bias into *gnd.* As the downstream transcriptional unit of *gnd* are O-antigen related genes (i.e. biosynthesis and export), this in turn lends further credence to a transcriptional regulatory model where biosynthesis of O-antigen and capsule is coupled for biophysical reasons as has been previously reported (66, 120). Collectively, these data show that O-antigen biosynthesis genes (i.e *wabI* and *waaL*) play a contributing role in serum resistance whilst *hns* mutants were shown to be more resistant to serum killing than WT *K. pneumoniae* ECL8. However, the mechanism that underpins this resistance is still largely unknown and requires further characterization. Despite the previously described links between *hns* and the regulation of a selection of capsular genes (66, 99), this is the first report of a role for *hns* in resisting complement-mediated killing in human serum.

This genome-wide screen has identified a suite of genetic fitness-factors required for growth of *K. pneumoniae* ECL8 in human urine and serum. The defined ECL8 urinome and serum resistome gene sets from this study gene differ from other TraDIS studies, hinting that the genotype of different pathogenic lineages might only be a piece in understanding the underlying infection biology. Future studies like these but expanding to other isolates, and the lesser studied clinically relevant *K. pneumoniae* species complex (KpSc), will be imperative for the effective future development of targeted therapeutics towards this problematic pathogen.

## Supporting information captions

**Figure S1:** (A) Pearson correlation coefficient (R2) of gene insertion index scores (IIS) of two sequenced technical replicates of the Klebsiella pneumoniae ECL8 TraDIS library (KTL1 and KTL2). The IIS of genes located on the chromosome and plasmid are highlighted in blue and red respectively. (B) Number of unique reads identified (%) in the raw fastQ file of the *K. pneumoniae* TraDIS library in sequentially larger k-mer pools up to three million reads. Plot generated using BBTools: bbcountunique.sh in non-cumulative mode (https://jgi.doe.gov/data-and-tools/bbtools/)

**Figure S2** Sequencing depth of the *K. pneumoniae* ECL8 plasmid (black) relative to the *K. pneumoniae* ECL8 chromosome (green). The plasmid has a read depth 1.33x that of the genome suggesting the plasmid has a copy number of one. Figure generated using Bandage (v3).

**Figure S3** The frequency distribution of insertion index scores. The insertion index score for each coding sequence was calculated as the number of insertions per CDS divided by the CDS length in base pairs to normalize for gene length. An exponential distribution model was fitted to the left mode that includes essential genes, and a gamma distribution model was fitted to the right, nonessential mode (blue). For a given insertion index score, the probability of belonging to each mode was calculated, and the ratio of these values was the log likelihood score. A gene was classified as essential if its log likelihood score was less than log2 12 and was therefore 12 times more likely to belong to the red mode than the blue mode.

**Figure S4** The COG Enrichment index comprising 373 genes classified as essential in *K. pneumoniae* ECL8. This index is calculated as the percentage of the essential genome made up of a COG divided by the percentage of the whole genome made up by the same COG. The log^2^ fold enrichment is displayed, and significant differences were calculated using the two-tailed Fisher’s exact test. Annotations were computed using eggnog-mapper based on eggNOG orthology data (2)

**Figure S5** Mathematical simulation (10^5^ instances) of random transposon insertion events under the null model of random insertion previously described (3). The probability of at least one insertion free region (IFR) of length (l) occurring in a genome of 5.3 Mb containing 554,834 transposon insertions (blue). Genome length and no. of genome-wide insertions from TraDIS studies by Langridge *et al.,* 2012 and Goodall *et al.,* 2018 plotted for comparison (3, 4).

**Figure S6 Sequencing depth required to sample a given proportion of the K. pneumoniae TraDIS library** (A) The following equation: 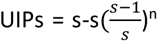 was applied to calculate the approximate number of sequence reads required to sample full library diversity i.e. 100% of Unique Insertion Points (UIPs). UIPs = Unique Insertion Points, n = number of mapped reads in millions (M) and s = sample size i.e. 499,919 CDS UIPs (B) The approximate number of CDS UIPs represented by a given number of sequence reads (M) are shown as a percentage of the total CDS UIPs i.e. 499,919.

**Figure S7** The Pearson correlation coefficient (R^2^) of gene insertion index scores (IIS) for two sequenced biological replicates of the *K. pneumoniae* ECL8 TraDIS library following thee 12 h passages in (blue) LB broth or (red) pooled human urine. The inline barcode identifiers used to demultiplex and distinguish replicates are highlighted in brackets ().

**Figure S8** Serum killing assay of *K. pneumoniae* ECL8 and *E. coli* BW25113. A sample of 2×10^8^ bacterial cells were incubated in 100 μL of human serum (solid line) or heat-inactivated human serum (dashed line) for 180 min. Viable bacterial numbers (cfu/ml) were sampled at regular time points by plating onto LB agar, overnight incubation at 37 °C and subsequent counting of the colonies. The log10 fold change in viable cell number was measured. *E. coli* BW25113 lacks O-antigen and was used as a serum sensitive control. The mean of three biological replicates is shown ± 1 SD.

**Figure S9** The pearson correlation coefficient (R^2^) of gene insertion index scores (IIS) for two sequenced biological replicates of the *K. pneumoniae* ECL8 TraDIS library following 90 min exposure to (blue) Heat-inactivated serum or (red) Serum. The inline barcodes used to demultiplex and distinguish replicates are given in brackets ().

**Figure S10** Pearson correlation coefficient (R2) of two biological replicates from the output TraDIS library following exposure to human serum for 180 min.

**Figure S11** Serum killing assay of WT K. pneumoniae ECL8, hns7::Tn5 and hns18::Tn5. Overnight cultures of bacterial cells were incubated in a 1:1 ratio (2×10^8^ cells : human serum) in a final volume of 100 μL. Viable bacterial numbers (CFU/mL) at 360 min timepoint was calculated by plating onto LB agar, overnight incubation at 37°C and subsequent counting of the colonies with a PBS control performed in parallel. The mean of three biological replicates is shown ± 1 SD

**Figure S12** (A) Transposon and read count insertion profiles of hns locus: red illustrating pooled mutant serum exposed for 180 min and blue denoting the before serum exposure input control. (B) Directional insertion bias of transposon (Tn) into *gnd*. Transposon insertions configured in the forward orientation (green), reverse orientation (blue). Transposon insertion densities are capped at a maximum read depth of 2000. Below – genomic context of *gnd* with a putative promoter for *manC* driving the transcriptional unit (*rfb* cluster) for O-antigen biosynthesis.

**Dataset S1** Essential gene table ECL8

**Dataset S2** Insertion Free Regions (IFR) within ECL8 (LB)

**Dataset S3** Dataset S3_Essential Gene Comparison against ECL8

**Dataset S4** Essential gene table ECL8 (Urine)

**Dataset S5** Essential gene table ECL8 (Serum)

**Dataset S6** Kaptive webserver results - ECL8 O-antigen K-antigen

**Dataset S7** AlbaTraDIS ECL8 180 Serum exposure results

