## Supplementary Datasets, Figures and Tables for "Transposon mutagenesis screen in *Klebsiella pneumoniae* identifies genetic determinants required for growth in human urine and serum": 2024 Feb - resubmission Gray et al. KP ECL8 TraDIS SUPP Figures and Tables.docx

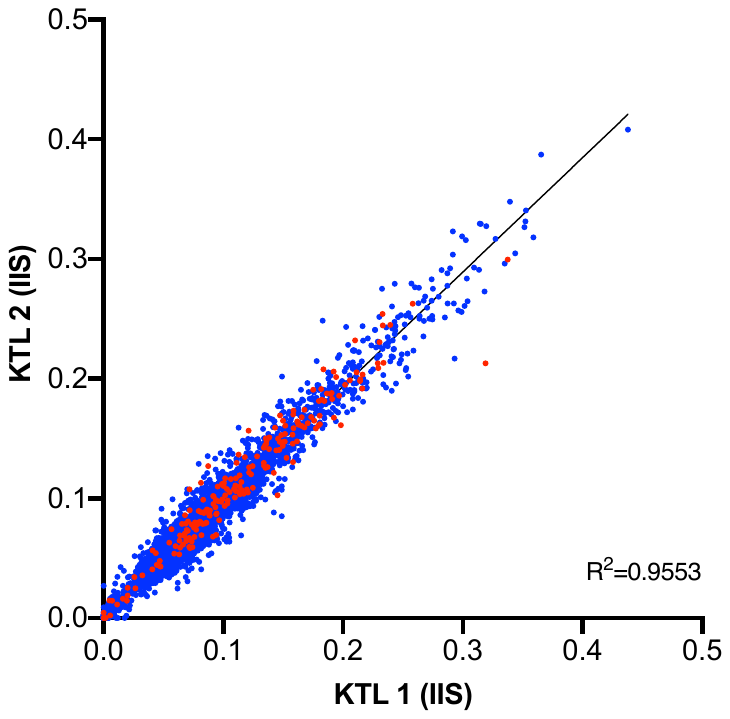

**B**

**A**

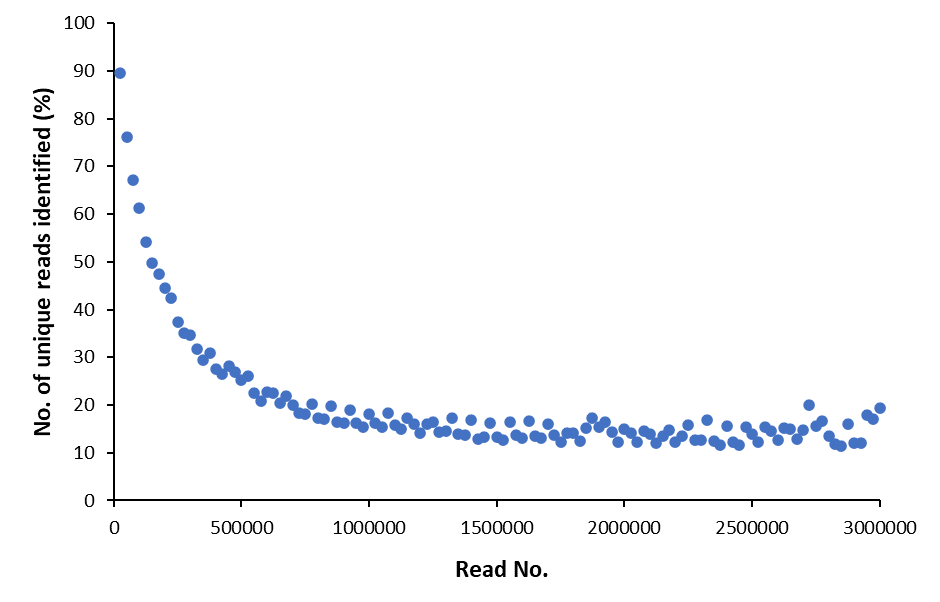

**Figure S1:** (A) Pearson correlation coefficient (R2) of gene insertion index scores (IIS) of two sequenced technical replicates of the Klebsiella pneumoniae ECL8 TraDIS library (KTL1 and KTL2). The IIS of genes located on the chromosome and plasmid are highlighted in blue and red respectively. (B) Number of unique reads identified (%) in the raw fastQ file of the *K. pneumoniae* TraDIS library in sequentially larger k-mer pools up to three million reads. Plot generated using BBTools: bbcountunique.sh in non-cumulative mode (<https://jgi.doe.gov/data-and-tools/bbtools/>)

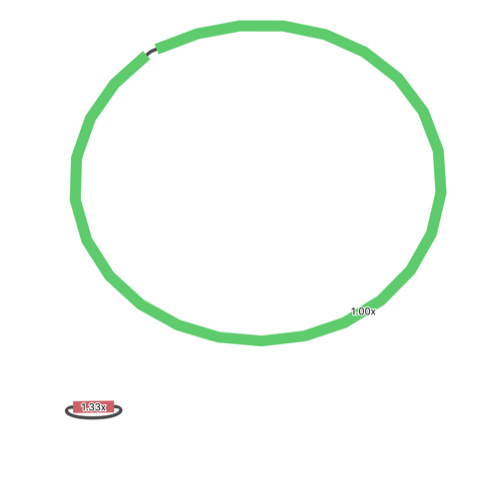
**Figure S2** Sequencing depth of the *K. pneumoniae* ECL8 plasmid (black) relative to the *K. pneumoniae* ECL8 chromosome (green). The plasmid has a read depth 1.33x that of the genome suggesting the plasmid has a copy number of one. Figure generated using Bandage (v3) (1).

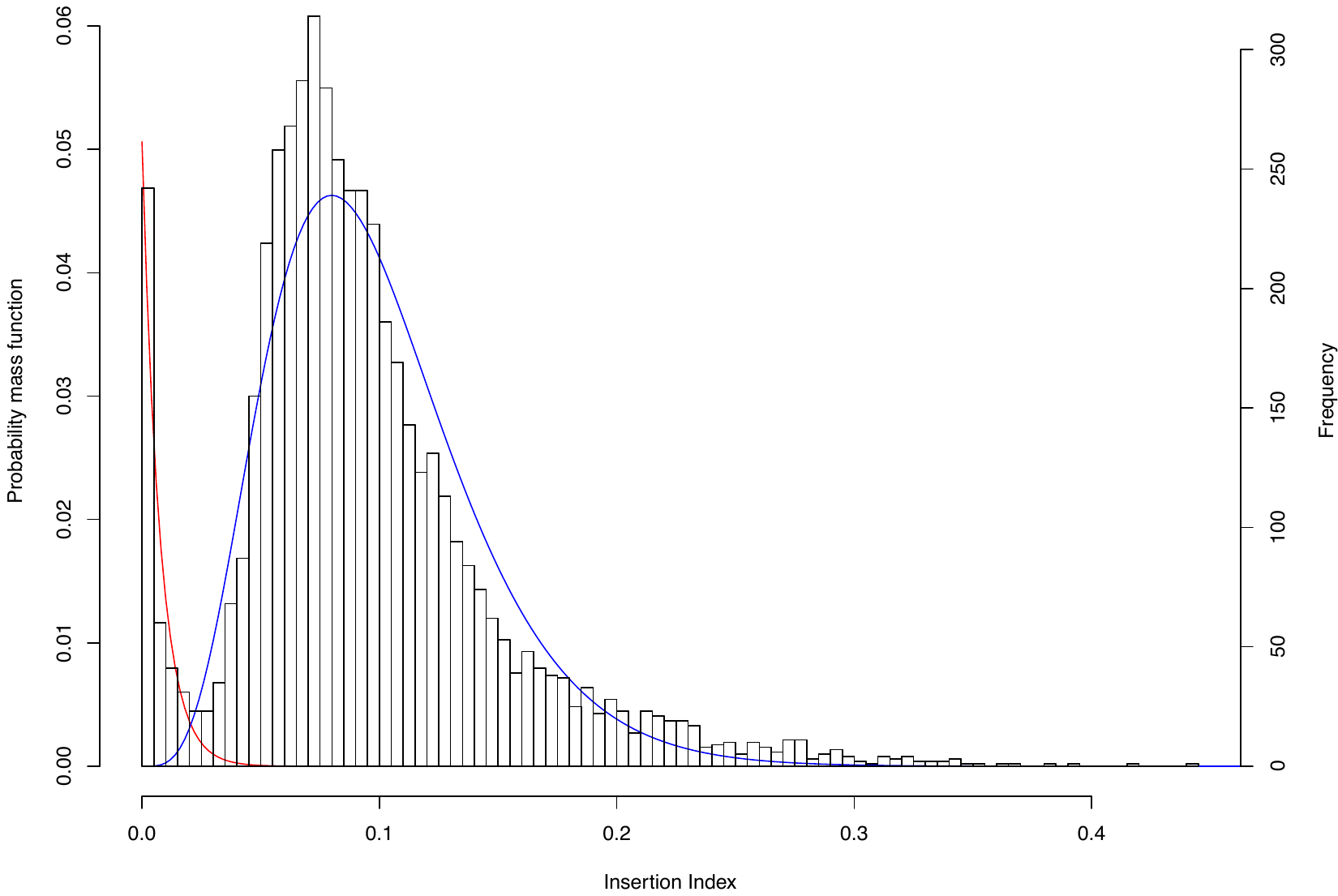

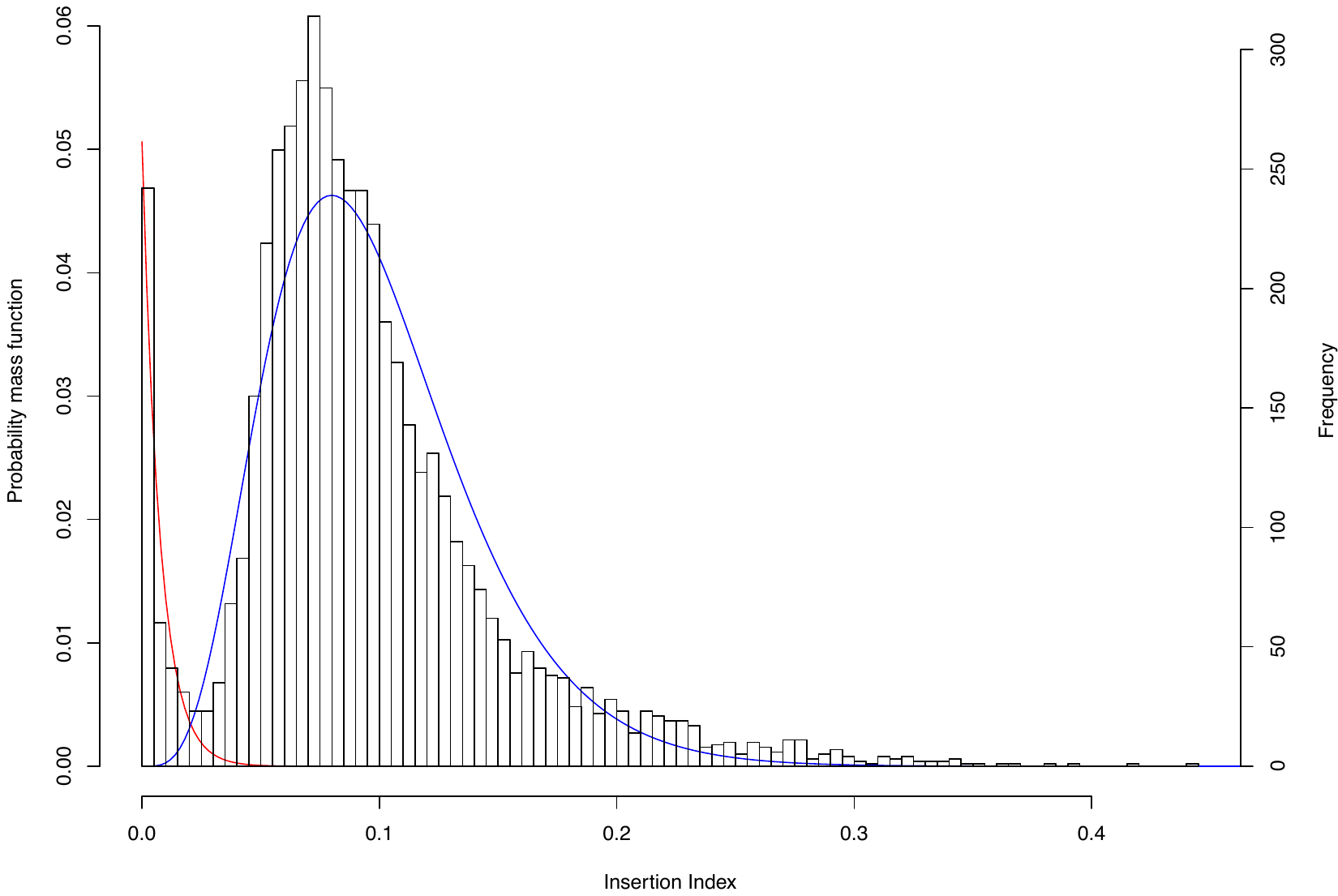
**Figure S3** The frequency distribution of insertion index scores. The insertion index score for each coding sequence was calculated as the number of insertions per CDS divided by the CDS length in base pairs to normalize for gene length. An exponential distribution model was fitted to the left mode that includes essential genes, and a gamma distribution model was fitted to the right, nonessential mode (blue). For a given insertion index score, the probability of belonging to each mode was calculated, and the ratio of these values was the log likelihood score. A gene was classified as essential if its log likelihood score was less than log_2_ 12 and was therefore 12 times more likely to belong to the red mode than the blue mode.

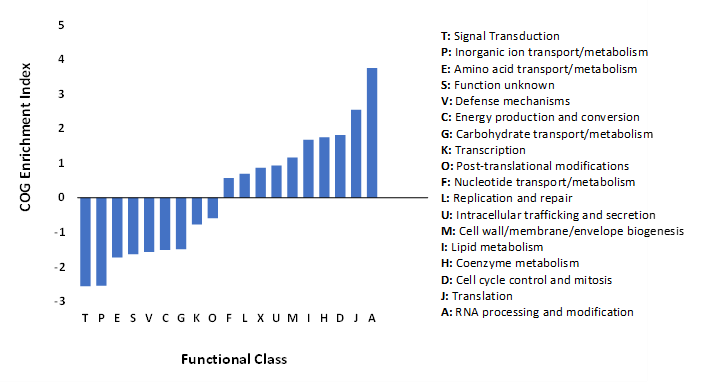
**Figure S4** The COG Enrichment index comprising 373 genes classified as essential in *K. pneumoniae* ECL8. This index is calculated as the percentage of the essential genome made up of a COG divided by the percentage of the whole genome made up by the same COG. The log^2^ fold enrichment is displayed, and significant differences were calculated using the two-tailed Fisher's exact test. Annotations were computed using eggnog-mapper based on eggNOG orthology data (2)

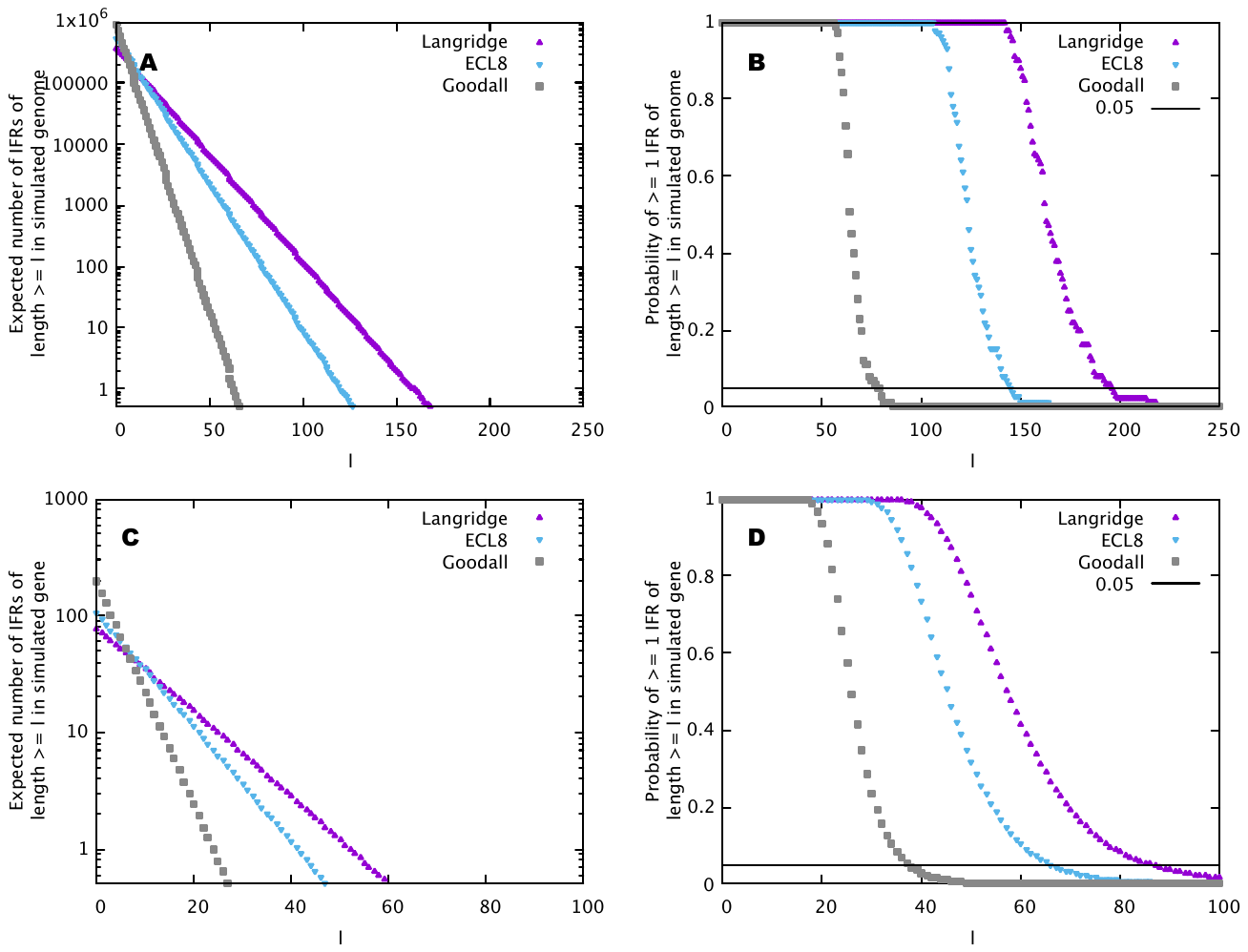

**Figure S5** Mathematical simulation (10^5^ instances) of random transposon insertion events under the null model of random insertion previously described (3). The probability of at least one insertion free region (IFR) of length (l) occurring in a genome of 5.3 Mb containing 554,834 transposon insertions (blue). Genome length and no. of genome-wide insertions from TraDIS studies by Langridge *et al.,* 2012 and Goodall *et al.,* 2018 plotted for comparison (3, 4).

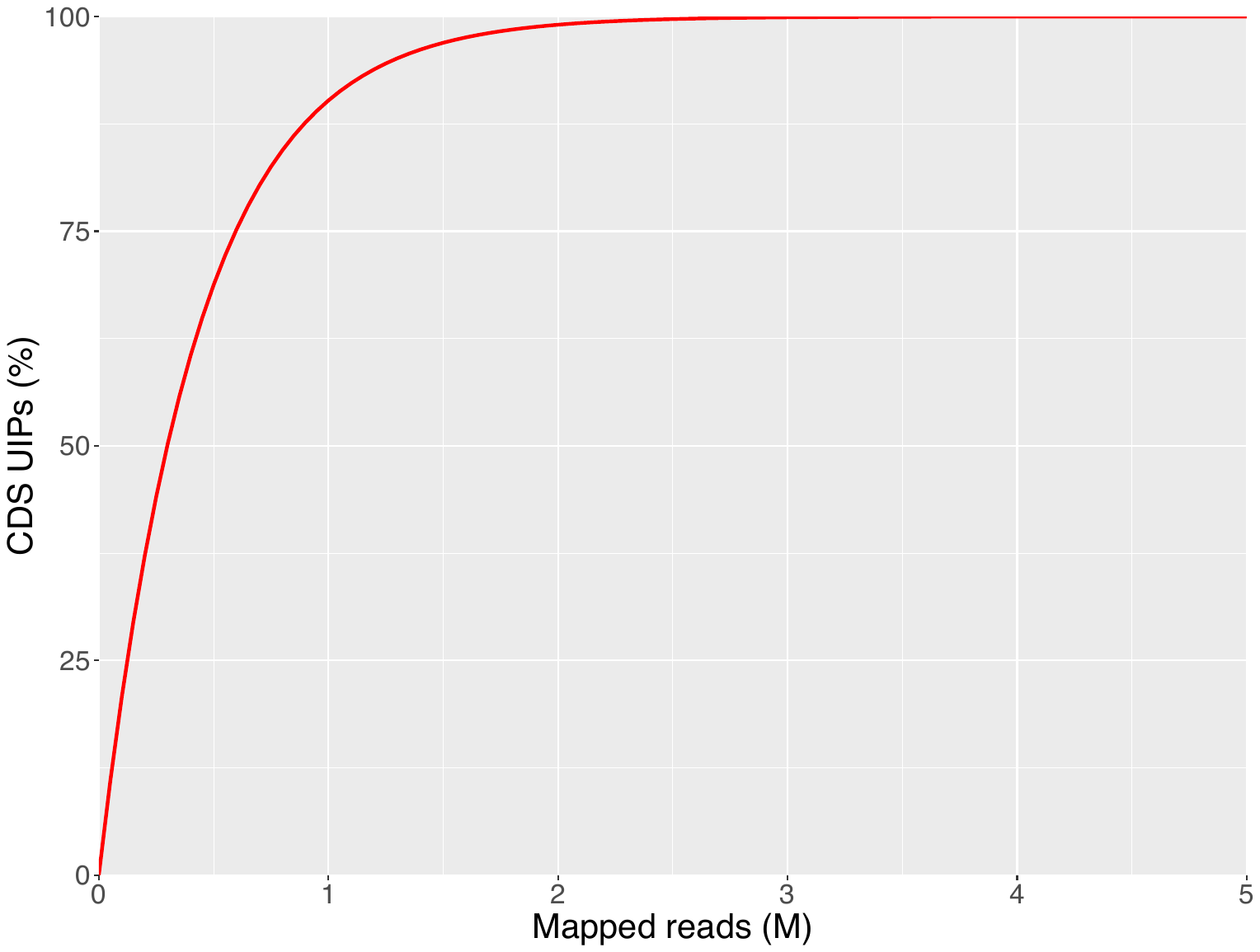

**A**

**B**

| **Number of reads (M)** | **CDS UIPs** | **% of total CDS UIPs** |
| --- | --- | --- |
| 0.5 | ~316,057 | 63.22 |
| 1 | ~432,284 | 84.47 |
| 2 | ~490,768 | 98.17 |
| 3 | ~498,681 | 99.75 |
| 4 | ~499,751 | 99.96 |

**Figure S6 Sequencing depth required to sample a given proportion of the K. pneumoniae TraDIS library** (A) The following equation: UIPs = s-s$(\frac{s-1}{s}$)^n^ was applied to calculate the approximate number of sequence reads required to sample full library diversity i.e. 100% of Unique Insertion Points (UIPs). UIPs = Unique Insertion Points, n = number of mapped reads in millions (M) and s = sample size i.e. 499,919 CDS UIPs (B) The approximate number of CDS UIPs represented by a given number of sequence reads (M) are shown as a percentage of the total CDS UIPs i.e. 499,919.

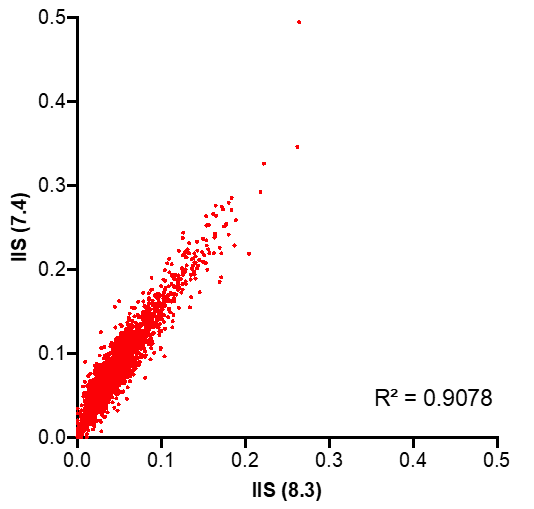

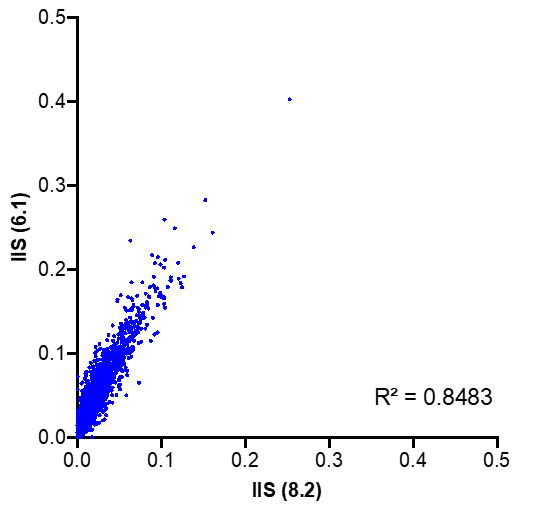

**Figure S7** The Pearson correlation coefficient (R^2^) of gene insertion index scores (IIS) for two sequenced biological replicates of the *K. pneumoniae* ECL8 TraDIS library following thee 12 h passages in (blue) LB broth or (red) pooled human urine. The inline barcode identifiers used to demultiplex and distinguish replicates are highlighted in brackets ().

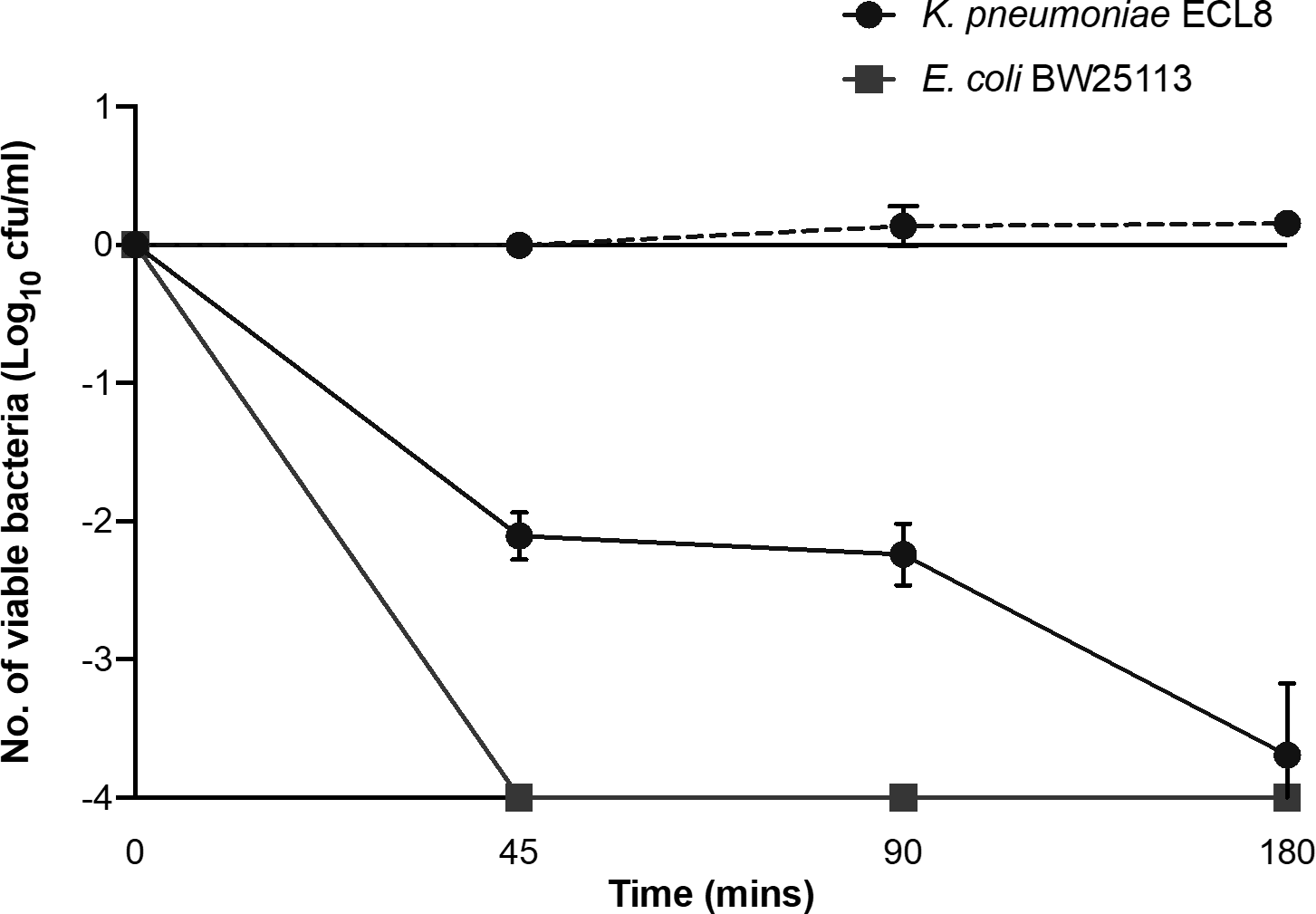

**Time (min)**

**Figure S8** Serum killing assay of *K. pneumoniae* ECL8 and *E. coli* BW25113. A sample of 2×10^8^ bacterial cells were incubated in 100 μL of human serum (solid line) or heat-inactivated human serum (dashed line) for 180 min. Viable bacterial numbers (cfu/ml) were sampled at regular time points by plating onto LB agar, overnight incubation at 37 °C and subsequent counting of the colonies. The log_10_ fold change in viable cell number was measured. *E. coli* BW25113 lacks O-antigen and was used as a serum sensitive control. The mean of three biological replicates is shown ± 1 SD.

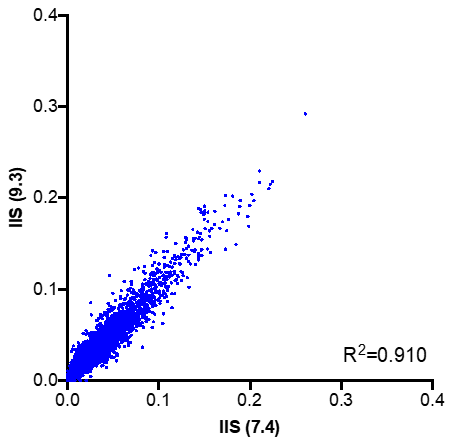

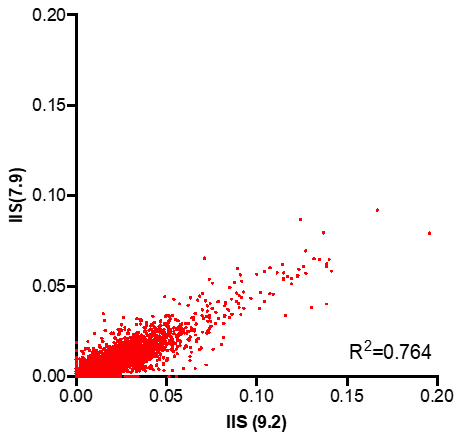
**F****i****gure S9** The pearson correlation coefficient (R^2^) of gene insertion index scores (IIS) for two sequenced biological replicates of the *K. pneumoniae* ECL8 TraDIS library following 90 min exposure to (blue) Heat-inactivated serum or (red) Serum. The inline barcodes used to demultiplex and distinguish replicates are given in brackets ().

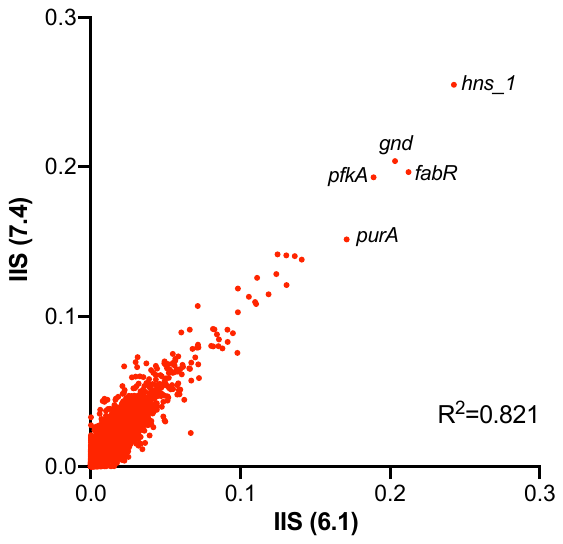

**Figure S10** Pearson correlation coefficient (R2) of two biological replicates from the output TraDIS library following exposure to human serum for 180 min.

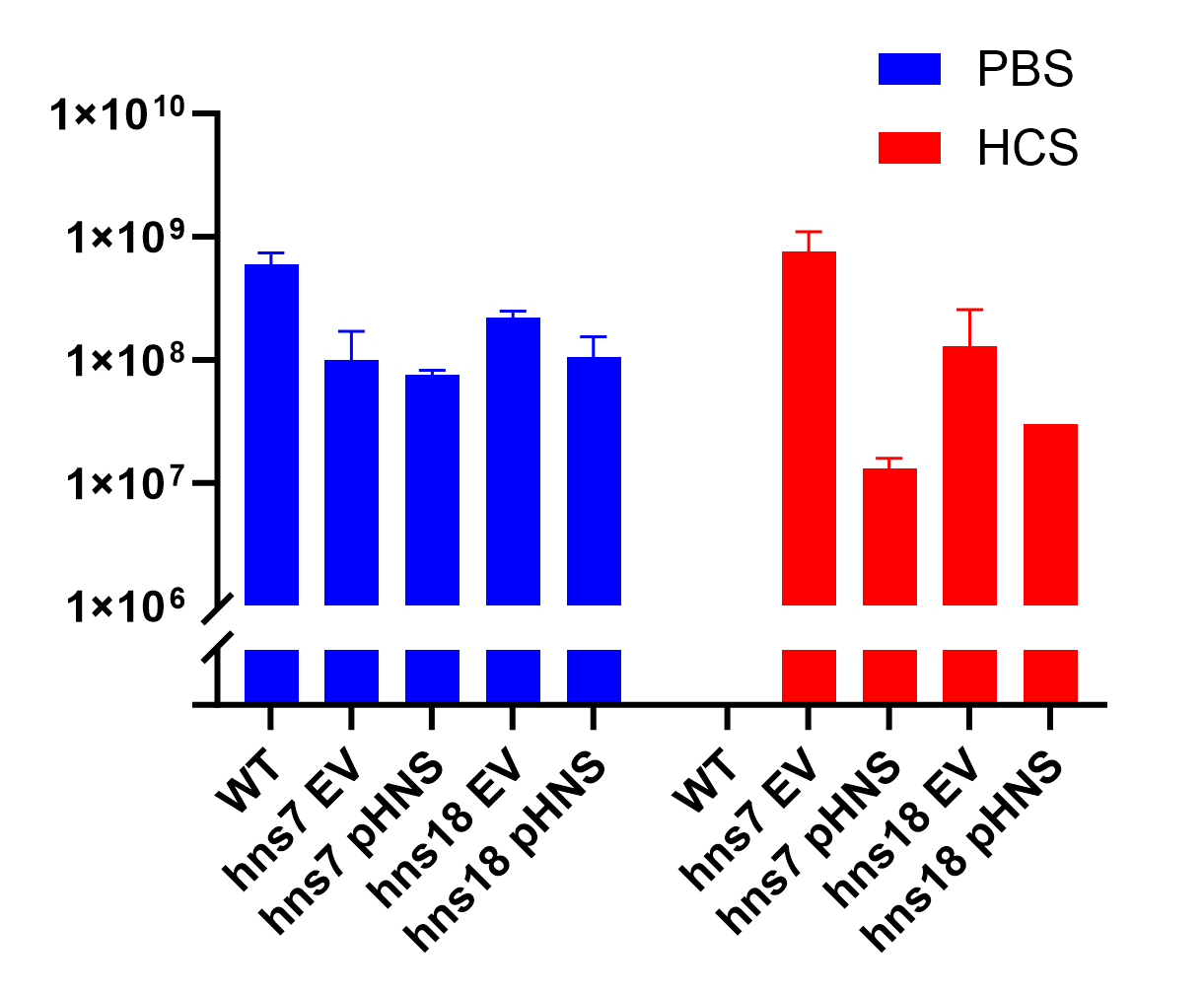

**Figure S11** Serum killing assay of WT K. pneumoniae ECL8, *hns*7::Tn5 and *hns*18::Tn5. Overnight cultures of bacterial cells were incubated in a 1:1 ratio (2×10^8^ cells : human serum) in a final volume of 100 μL. Viable bacterial numbers (CFU/mL) at 360 min timepoint was calculated by plating onto LB agar, overnight incubation at 37°C and subsequent counting of the colonies with a PBS control performed in parallel. The mean of three biological replicates is shown ± 1 SD

**A**

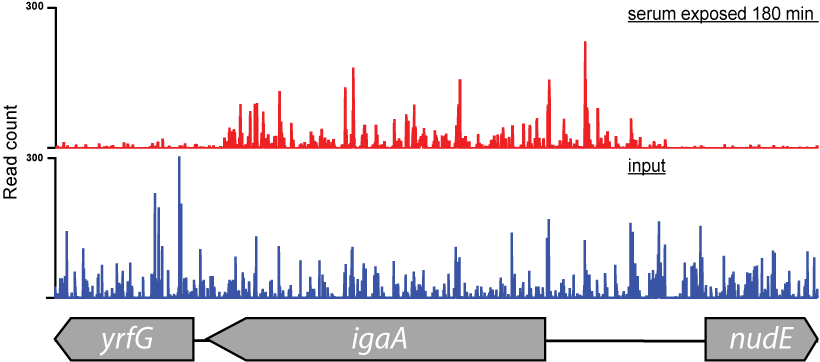

**B**

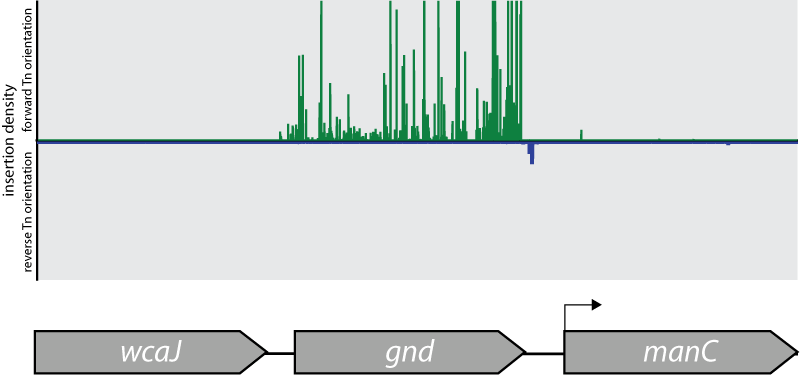

**Figure S12** (A) Transposon and read count insertion profiles of hns locus: red illustrating pooled mutant serum exposed for 180 min and blue denoting the before serum exposure input control. (B) Directional insertion bias of transposon (Tn) into *gnd*. Transposon insertions configured in the forward orientation (green), reverse orientation (blue). Transposon insertion densities are capped at a maximum read depth of 2000. Below – genomic context of *gnd* with a putative promoter for *manC* driving the transcriptional unit (*rfb* cluster) for O-antigen biosynthesis.

**Table S1: Bacterial strains and plasmids utilised in this study**

|  | **Description** | **Source** |
| --- | --- | --- |
| Strain |  |  |
| *K. pneumoniae* ECL8 | *K. pneumoniae* isolate derived from NCTC 00418 (Strep^R^) (Amp^R^) (Rif^R^) | (5) |
| *K. pneumoniae* ECL8 *sodA::aph* | *K. pneumoniae* ECL8 with the *sodA* gene replaced with a kanamycin *aph* cassette (Kan^R^) | This study |
| *K. pneumoniae* ECL8 *ytfL::aph* | *K. pneumoniae* ECL8 with the *ytfL* gene replaced with a kanamycin *aph* cassette (Kan^R^) | This study |
| *K. pneumoniae* ECL8 *ompA::aph* | *K. pneumoniae* ECL8 with the *ompA* gene replaced with a kanamycin *aph* cassette (Kan^R^) | This study |
| *K. pneumoniae* ECL8 *fepB::aph* | *K. pneumoniae* ECL8 with the *fepB* gene replaced with a kanamycin *aph* cassette (Kan^R^) | This study |
| *K. pneumoniae* ECL8 *fepD::aph* | *K. pneumoniae* ECL8 with the *fepD* gene replaced with a kanamycin *aph* cassette (Kan^R^) | This study |
| *K. pneumoniae* ECL8 *exbB::aph* | *K. pneumoniae* ECL8 with the *exbB* gene replaced with a kanamycin *aph* cassette (Kan^R^) | This study |
| *K. pneumoniae* ECL8 *exbD::aph* | *K. pneumoniae* ECL8 with the *exbD* gene replaced with a kanamycin *aph* cassette (Kan^R^) | This study |
| *K. pneumoniae* ECL8 *wbbY::aph* | *K. pneumoniae* ECL8 with the *wbbY* gene replaced with a kanamycin *aph* cassette (Kan^R^) | This study |
| plasmid |  |  |
| pKD4 | Used as a template for amplification of the kanamycin aph cassette for the construction of chromosomal mutations. | (6) |
| pACBSCE | Arabinose inducible plasmid that encodes for λ-Red genes: gam, exo and bet llelic exchange vector with arabinose induction | (7) |

**Table S2: Primer nucleotide sequences for construction of *K. pneumoniae* chromosomal mutant strains.**

| **Name** | **Primer sequence (5’-3’)** | **Description*** |
| --- | --- | --- |
| ompA_F | GCTAAACTGGGTTACCCGATCACTGACGATCTGGACATCTACACCCG  TCTGGGCGGCATGGTGTAGGCTGGAGCTGCTTC | Forward primer for replacement of *ompA* with the kanamycin *aph* cassette |
| ompA_R | GGGCATAAAAAAAACCCGCCGAAGCGGGTTTTTTTTTATCGGTTATA  ACTTAAGCCGCCGGCTGAGTTACCATATGAATATCCTCCTTAG | Reverse primer for replacement of *ompA* with the kanamycin *aph* cassette |
| ompA_checkF | CAGCTTGGTGCTGGTGCGTTC | Forward check primer for confirmation of the correct disruption of *ompA* |
| ompA_checkR | GCAAAATGTCGAGCCTGAAG | Reverse check primer for confirmation of correct disruption of *ompA* |
| sodA_F | GGACAAACTGCTTACGCGGCGTTAACACTTGAGCCGCTCGACAATAA  TGGAGATGATTATGGTGTAGGCTGGAGCTGTTC | Forward primer for replacement of *sodA* with the kanamycin *aph* cassette |
| sodA_R | GCGAGTCATCAGACTCGCTTCTCTTTTAGCGCAATGCAACCTTATTTTT  TGGCGGCAAAACGCATATGAATATCCTCCTTAG | Reverse primer for replacement of *sodA* with the kanamycin *aph* cassette |
| sodA_checkF | TGCCACAGGATGGCGAACCC | Forward check primer for confirmation of the correct disruption of *sodA* |
| sodA_checkR | GGTCCCCATCGAGACCGAGA | Reverse check primer for confirmation of correct disruption of *sodA* |
| exbB_F | TGTCGTTTTGATATTATTGTGGGCAGATTTTGTGATTATCGTCGTGGAG  ATAGAGCGTGGTGTAGGCTGGAGCTGCTTC | Forward primer for replacement of *exbB* with the kanamycin *aph* cassette |
| exbB_R | CGCCGTTATCGTCCAGGTTTTCATTAAGACGCATCGCCATAGCCGATCA  ACCTACCCGCATATGAATATCCTCCTTAG | Reverse primer for replacement of *exbB* with the kanamycin *aph* cassette |
| exbB_checkF | GCTTTTCTATACCAGCGCACCG | Forward check primer for confirmation of the correct disruption of *exbB* |
| exbB_checkR | GATGGGTTTTTCCGGTCGCGGT | Reverse check primer for confirmation of correct disruption of *exbB* |
| exbD_F | CCAGCGGCGTGAAGCCGGTGCGCAGCGCGCAGAAATTACGGGTAG  GTTGATCGGCTATGGTGTAGGCTGGAGCTGCTTC | Forward primer for replacement of *exbD* with the kanamycin *aph* cassette |
| exbD_R | CAACAAAAAAAGGCCTGCACGCGGCCAGCCTTTGCAGAAACGCAAG  CGGGTTATTTGGCCATATGAATATCCTCCTTAG | Reverse primer for replacement of *exbD* with the kanamycin *aph* cassette |
| exbD_checkF | GTCCTCTGATTCTACGAGGCACG | Forward check primer for confirmation of the correct disruption of *exbD* |
| exbD_checkR | AAATAAACCGGCGCCAGCAGCC | Reverse check primer for confirmation of correct disruption of *exbD* |
| ytfL_F | CACATTTGAGTTATCAACTTCCCTTCCGAGGATCTGGCCTCAACGGT  CAGAAAAGATATGGTGTAGGCTGGAGCTGCTTC | Forward primer for replacement of *ytfL* with the kanamycin *aph* cassette |
| ytfL_R | GGCGGATGGTCATCCGCCCTTAGGAGAGAGAAAAGATTACGCTCAG  GCGTTCTGGCTTTCCATATGAATATCCTCCTTAG | Reverse primer for replacement of *ytfL* with the kanamycin *aph* cassette |
| ytfL_checkF | CTAGCCAGTGTGACAGCCGG | Forward check primer for confirmation of the correct disruption of *ytfL* |
| ytfL_checkR | GCTATCGGCAGAGGGGCGTG | Reverse check primer for confirmation of correct disruption of *ytfL* |
| fepB_F | CACAAAGTTGAAAATGAGACGCATTTATCACCTTTCAAATCAGGAT  GCGATGACGTGGTGTAGGCTGGAGCTGCTTC | Forward primer for replacement of *fepB* with the kanamycin *aph* cassette |
| fepB_R | GCAGGCCGAGTGCCCGTCCTGATGGCGCAGCCCGCGTTAGCCGAAC  AGGCTGGAGAGCATATGAATATCCTCCTTAG | Reverse primer for replacement of *fepB* with the kanamycin *aph* cassette |
| fepB_checkF | TGACGTTTCCATATCATCCTC | Forward check primer for confirmation of the correct disruption of *fepB* |
| fepB_checkR | GCTGGCATTGTAGGCCGGGC | Reverse check primer for confirmation of correct disruption of *fepB* |
| fepD_F | TGAATAAAATCGATAACGATAATTACTATCATTATCATATCAGGGATG  TCAGTTATGGTGTAGGCTGGAGCTGCTTC | Forward primer for replacement of *fepD* with the kanamycin *aph* cassette |
| fepD_R | GCAGACAGCTGGCTATCAGGCGGCGGGACGGGGCAATCACAGGCCA  CCTCCCCGCGGCATATGAATATCCTCCTTAG | Reverse primer for replacement of *fepD* with the kanamycin *aph* cassette |
| fepD_checkF | GCGAGCGATAAAAACGGCGC | Forward check primer for confirmation of the correct disruption of *fepD* |
| fepD_checkR | CCATTAACACCCGCGGCAGC | Reverse check primer for confirmation of correct disruption of *fepD* |
| wbbY_F | ACTACTTCAATTCACTAATATCATAGAAAAGTCTAGGTTACAAAGGA  AGGGTTACAATGGTGTAGGCTGGAGCTGCTTC | Forward primer for replacement of *wbbY* with the kanamycin *aph* cassette |
| wbbY_R | GAAAGTTAATATTGTTTTTGCGGAGCCCTTTCGGGCCCCGAATATTA  CTTTATTTTAACCATATGAATATCCTCCTTAG | Reverse primer for replacement of *wbbY* with the kanamycin *aph* cassette |
| wbbY_checkF | TTACACCATCACCAGCATTAC | Forward check primer for confirmation of the correct disruption of *wbbY* |
| wbbY_checkR | TCCGGCTGAATTCATCCGAAG | Reverse check primer for confirmation of correct disruption of *wbbY* |

*Check primers annealing ~200 bp upstream and downstream of the gene of interest were utilised for confirmation of mutant strains by PCR and sanger sequencing

**Table S3:** **Primer nucleotide sequences for enrichment of the transposon junction (TKK_F and TKK_R) and the introduction of an inline barcode for multiplexed sequencing (TKK 6, 7, 8, 9)**

| **Name** | **Primer sequence (5’-3’) *** |
| --- | --- |
| TKK 6.1 | AATGATACGGCGACCACCGAGATCTACACTCTTTCCCTACACGACGCTCTTCCGATCTCGTACGAGCTTCAGGGTTGAGATGTGTA |
| TKK 6.3 | AATGATACGGCGACCACCGAGATCTACACTCTTTCCCTACACGACGCTCTTCCGATCTTACGTAAGCTTCAGGGTTGAGATGTGTA |
| TKK 7.2 | AATGATACGGCGACCACCGAGATCTACACTCTTTCCCTACACGACGCTCTTCCGATCTGCTAGCTAGCTTCAGGGTTGAGATGTGTA |
| TKK 7.4 | AATGATACGGCGACCACCGAGATCTACACTCTTTCCCTACACGACGCTCTTCCGATCTTAGCTAGAGCTTCAGGGTTGAGATGTGTA |
| TKK 8.2 | AATGATACGGCGACCACCGAGATCTACACTCTTTCCCTACACGACGCTCTTCCGATCTATGCATGCAGCTTCAGGGTTGAGATGTGTA |
| TKK 8.3 | AATGATACGGCGACCACCGAGATCTACACTCTTTCCCTACACGACGCTCTTCCGATCTCATGCATGAGCTTCAGGGTTGAGATGTGTA |
| TKK 8.4 | AATGATACGGCGACCACCGAGATCTACACTCTTTCCCTACACGACGCTCTTCCGATCTCGTACGAGCTTCAGGGTTGAGATGTGTA |
| TKK 9.2 | AATGATACGGCGACCACCGAGATCTACACTCTTTCCCTACACGACGCTCTTCCGATCTATCGATCGAAGCTTCAGGGTTGAGATGTGTA |
| TKK 9.3 | AATGATACGGCGACCACCGAGATCTACACTCTTTCCCTACACGACGCTCTTCCGATCTTCGATCGATAGCTTCAGGGTTGAGATGTGTA |
| TKK 9.4 | AATGATACGGCGACCACCGAGATCTACACTCTTTCCCTACACGACGCTCTTCCGATCTCGATCGATCAGCTTCAGGGTTGAGATGTGTA |
| TKK_F | ACCTGCAGGCATGCAAGCTTCAGG |
| TKK_R | GACTGGAGTTCAGACGTGTGCTCTTCCGATC |

*The expected inline barcode is underlined
